# Spatiotemporal Regulatory Logics of Mouse Gastrulation

**DOI:** 10.1101/2024.12.22.630012

**Authors:** Xianfa Yang, Bingbing Xie, Penglei Shen, Yingying Chen, Chunjie Li, Fengxiang Tan, Yumeng Yang, Yun Yang, Rui Song, Panpan Mi, Zhiwen Liu, Mingzhu Wen, Patrick P. L. Tam, Shengbao Suo, Naihe Jing

## Abstract

Spatiotemporal coordination of cellular and molecular events is crucial for cell fate commitment during mouse gastrulation. However, the high-precision mechanisms governing the timing and spatial dynamics remain poorly understood. Here, we present a time-series single-cell multi-omic dataset from the mouse gastrulating embryos and construct a hierarchical gene regulatory landscape. Integrating this with real three-dimensional transcriptomic coordinate, we created ST-MAGIC and ST-MAGIC (+) atlas, dissecting the spatiotemporal logics of regulatory networks and signaling responsiveness underpinning the lineage commitment at gastrulation. Specifically, we delineated the multi-omic basis for left-right symmetry breaking events in the gastrula and also revealed the spatiotemporal molecular relay for axial mesendoderm lineage, where early and intermediate transcription factors first open the chromatin regions and setup the responsiveness to signaling, followed by terminal factors to consolidate the transcriptomic architecture. In summary, our study presents a spatiotemporal regulatory logic framework of mouse gastrulation, that advances our understanding of mammalian embryogenesis.

## Introduction

Gastrulation is a pivotal phase in embryonic development, at which the multipotent epiblast is allocated to diverse tissue lineages of the three primary germ layers: ectoderm, mesoderm, and endoderm.^1^ This stage is also crucial for the spatial organization of the embryo in anterior-posterior, dorsal-ventral and left-right axes.^1^ Gastrulation therefore lays the blueprint of embryogenesis, making it a focal point for understanding the molecular and cellular mechanisms that govern embryonic patterning.

The dynamic nature of gastrulation involves rapid changes in cell composition, differentiation and proliferation, all of which are intricately regulated by a combination of epigenetic modifications, transcriptional networks, and intercellular communication. Recent advancements using limited cell or single-cell omics significantly enhance our understanding of the dynamic processes during gastrulation. These technologies have enabled high-resolution analysis of gene expression patterns and cellular heterogeneity within developing embryos, identify that embryogenesis during gastrulation is orchestrated by precise schedules of gene expression within individual cells, regulated by complex intrinsic gene regulatory networks (GRNs) and influenced by intercellular signaling pathways.^2–5^ These findings highlight the importance of both temporal and spatial regulation in cell fate and tissue patterning.

Despite these advances, a comprehensive characterization of the GRNs that govern the rapid cell fate commitment and spatial patterning during gastrulation remains elusive. Traditional single modality of single-cell omics often disrupts the spatial and temporal continuity of embryonic development, limiting our ability to fully understand the molecular mechanisms underlying embryo patterning. Spatiotemporal coordination of GRNs provides the developmental principles of lineage allocation and tissue patterning in the germ layers of the gastrulating embryo.

To address these challenges, we generated a high-resolution, spatiotemporally resolved single-cell multi-omic reference for the mouse gastrulating embryo. Our dataset includes transcriptomics and chromatin accessibility profiles, collected at 6-hour intervals across multiple stages of gastrulation. Using the developed Bi-Orientation Cis-Regulatory Elements Predictor (Bio-CRE) algorithm, we linked genes with their potential regulatory elements, establishing a comprehensive GRNs atlas underlying the rapid cell lineage commitment process. With reference to the three-dimensional spatial transcriptome coordinates, the dataset is rendered into the SpatioTemporal-Multiomic Atlas of Gastrulating In-silico Cells (ST-MAGIC). By integrating transcription factor (TF)-target gene-target chromatin region cascades as well as signaling effector chromatin immunoprecipitation sequencing (ChIP-seq) data, we developed the ST-MAGIC (+) platform to explore the spatiotemporal dynamics of TF networks and determine intrinsic responsiveness to crucial developmental signals.

From the ST-MAGIC and ST-MAGIC (+) atlas, we have determined the multi-omic basis for left-right symmetry breaking events in the gastrula, identified and experimentally validated the complex GRNs hierarchy involving TFs (EOMES, FOXA2, NOTO/POU6F1) and signaling pathways (such as NODAL and WNT signaling) that regulate mouse axial mesendoderm development, accompanied by a spatial developmental route from the anterior primitive streak to the distal tip of the embryo.

## Results

### Single-cell multi-omics capture the GRNs underlying germ layer formation

To spatiotemporally dissect the intrinsic GRNs at single-cell resolution during mouse gastrulation, we have undertaken: (1) profiling the transcriptome and chromatin accessibility of individual cells of mouse gastrula at early-streak stage (E6.5) to the late-streak stage (E7.5) at 6-hour (0.25 embryonic day) intervals; (2) constructing gene-peak linkages, that connect transcriptomic and epigenomic modalities; (3) integrating the inferred gene-peak linkages with the Geo-seq-based spatiotemporal information to reconstruct a real time-series three-dimensional spatiotemporal multiome atlas; and (4) analyzing the hierarchical logics of GRNs in parallel with signaling responsiveness to gain insights into the mechanistic attributes of the gene regulatory networks underpinning lineage development and embryo patterning at gastrulation (Fig. 1a).

**Fig. 1.**
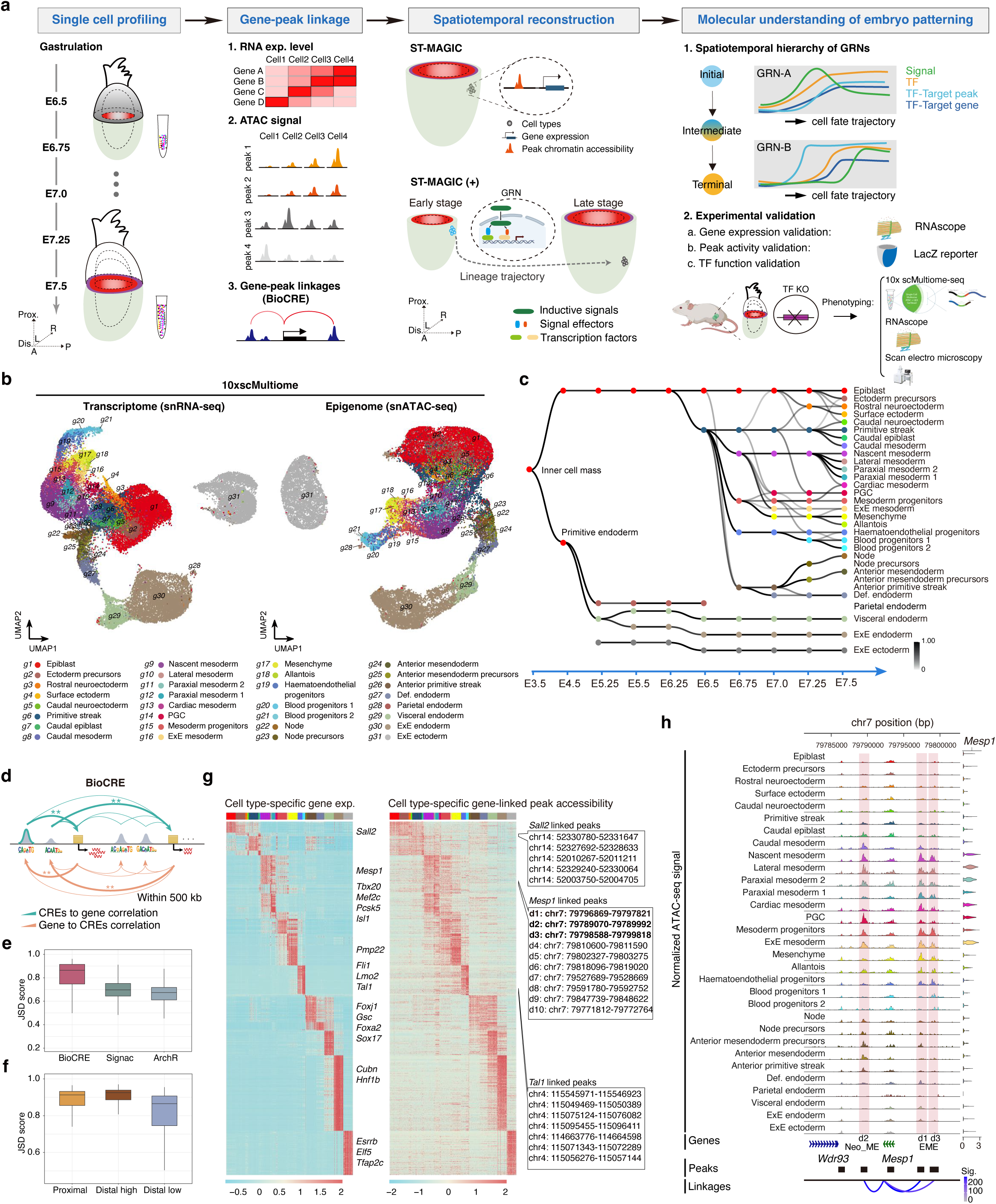
Building the single-cell multi-omic atlas of the mouse gastrulating embryo. **a.** Schematic overview of the work flow from data generation, peak-to-gene linkage identification to construction of spatiotemporal multiomic atlas spanning from E6.5 to E7.5 of mouse embryo development, followed by the determination of lineage-specific GRNs and experimental validation. **b.** UMAP projection of cells based on multiome-derived transcriptomic (left) and epigenomic (right) data colored by celltype annotations. The annotations of celltypes were inferred from the published dataset.^2,5^ **c.** Directed acyclic graph showing inferred relationships between celltypes defined by transcriptomic data across early mouse embryogenesis. The processed scRNA-seq dataset covering the developmental stages from E3.5 to E6.25 was publicly available from the TOME portal (http://tome.gs.washington.edu/). Each row corresponds to one of the annotated celltypes, columns to developmental stages. All edges with weights above 0.2 were shown in grayscale. Of note, parietal endoderm cells were not actively retained during the isolation of embryos in this study and were excluded from analyses in this study. **d.** The schematic diagram showing the capture of gene-peak linkages using BioCRE. **e.** The Jensen-Shannon divergence (JSD) index evaluating the performance for BioCRE, Signac, and ArchR algorithm in calculating the consistency between celltype transcriptomic features and chromatin accessibility. **f.** BioCRE specifically captures a group of distal peaks with the highest consistency with linked gene expression. Distal peaks were annotated as peaks located outside TSS ± 2 kb, proximal peaks were annotated as peaks located within TSS ± 2 kb. **g.** Heatmap showing gene expression (left) and chromatin accessibility (right) for selected celltype specific genes (top 10 genes) and their linked peaks with each row representing a pair of one gene and the linked CRE. Each gene can be linked to multiple CREs. Hence, genes can be represented by multiple rows in the corresponding heatmap. **h.** Genome browser snapshot displaying the normalized ATAC-seq signal (top), the distribution of annotated peaks (middle), the linkage with *Mesp1* gene for a given celltype (bottom). The expression of *Mesp1* across 31 celltypes was also shown at the right. Three genomic peaks showing significant linkages with *Mesp1* were highlighted in red, two of the peaks (d1, d3) consist the known EME enhancer reported previously^8^, and d2 peak was a newly identified *Mesp1*-linked element and named as Neo_ME element,. Sig. represents the significance of the linkage correlation between gene and chromatin peak calculated by BioCRE, the same for the following coverage plot results. For detailed significance calculation, please see the Method section.

In total, we have acquired bi-modal omics data from 35,449 single cells across five embryonic stages (E6.5, E6.75, E7.0, E7.25, and E7.5) with high cell coverage and high data quality after stringent quality control measurements (Extended Data Fig.1a-d). Uniform manifold approximation and projection (UMAP) was performed for both transcriptomics and chromatin accessibility (Fig. 1b; Extended Data Fig. 1e, 1f). To annotate the celltypes, iterative transcriptomic clustering for cells from each stage was conducted and specific gene expression pattern for each sub-cluster was refined. Cells with similar gene expression profiles were then grouped and annotated by inferring known gene expression signatures cross-referenced with the published mouse embryo transcriptome atlases.^2,5,6^ The 31 annotated celltypes exhibit consistent composition and high transcriptomic correlation with the published single-cell transcriptome gastrulation atlas^2^ and delineate the celltype composition diversifying process during gastrulation with some celltypes were selectively enriched for earlier (e.g. anterior primitive streak) or later stages (e.g. cardiac mesoderm etc.) (Extended Data Fig. 1g, 2a-d). Notably, in contrast to one homogeneous notochord cell cluster recognized in the previous report,^2^ here we identified four previously unappreciated distinct subgroups within the notochord lineage, which was named as anterior mesendoderm precursors and node precursors at the E7.25 stage, and anterior mesendoderm and node at the E7.5 stage, respectively (Fig. 1b).

The annotated cell identities were then transferred to cells embedded in the snATAC-seq UMAP based on the in silico cell-to-cell matches bridged by the common cellular barcodes incorporated during the preparation of multi-omic sequencing library (Fig. 1b). Generally, the cells collated in this dataset cover the transitions from early pluripotent epiblast cells at E6.5 to fate-committed neuroectoderm, definitive endoderm and various diversified mesodermal subtypes at E7.5 stage. We also applied the Trajectories Of Mammalian Embryogenesis (TOME) algorithm^6^ to systematically analyze the relationship among celltypes from two adjacent stages and finally generated a clear relationship graph for all the celltypes across gastrulation (Fig. 1c). The inferred lineage trajectories are broadly consistent with previous understandings of mouse early embryogenesis,^2,6,7^ providing a developmental phylogeny to pinpoint the molecular dynamics and hierarchy of transcriptomic and epigenetic regulation during gastrulation.

To map the GRNs governing lineage development during gastrulation, we first determined celltype-specific gene expression, chromatin accessibility and motif enrichment. Intriguingly, despite distinct celltypes exhibit distinctive transcriptomic features and epigenomic differences, the motif enrichment remained intermingled for cells within the same pedigree (Extended Data Fig. 3a-c). This observation may indicate that the formation of various celltypes and the transcriptomic signatures may be regulated by a common set of TFs but with frequent turnover of epigenomic landmarks. Next, to capture the relationship between gene transcriptomic status and peak chromatin accessibility, we developed an algorithm, BioCRE, to capture the linkage between expressed genes and candidate regulatory elements, thereby gaining insight into the relationship between gene expression and chromatin peak accessibility. Distinct from existing tools, such as Signac and ArchR, BioCRE harnesses a bi-orientation regression model leveraging multi-omics data at chromosome level to identify potential CREs (Fig. 1d). The Jensen–Shannon Divergence (JSD) index, measuring the consistency between celltype transcriptomic features and linked chromatin accessibility diversities, was higher for BioCRE than for Signac and ArchR (Fig. 1e). This result indicates that BioCRE would be more effective in predicting celltype-specific regulatory elements that modulate target gene expression.

Generally, BioCRE results show that one gene is linked to a median of 5 CREs with a median gene-to-peak distance of 127,175 bp (Extended Data Fig. 3d-f). Moreover, we observed that a considerable number of distal chromatin peaks (TSS ± 2-500 kb) exhibit higher JSD index (distal high group) than the regulated gene promoters, highlighting the distal regulatory regions may play prominent roles in regulating gene expression (Fig. 1f). Co-variation of celltype-specific DEGs and BioCRE-linked peaks’ chromatin accessibilities faithfully distinguished the identified celltypes in both transcriptomic and epigenomic modalities (Fig. 1g; Supplementary Table 1). For example, ten distal peaks were linked to the expression of nascent mesoderm marker, *Mesp1* (Fig. 1g). Amongst the distal linked peaks, d1 and d3 peaks are related to the previously known EME enhancer for *Mesp1*,^8^ while the potential regulatory logics for the remaining eight distal peaks were newly identified. To validate the regulatory force for the newly-identified *Mesp1* linked distal peaks, we analyzed the chromatin accessibilities attribute for one of the newly-identified distal peaks (named as Neo_ME) and the EME element (Fig. 1h). Detailed exploration revealed that, apart from the co-accessible patterns in nascent mesoderm, lateral mesoderm, paraxial mesoderm 2, mesoderm progenitors and ExE mesoderm cells, the Neo-ME element was accessible in the caudal epiblast, caudal mesoderm cells as well as axial mesendoderm (node and anterior mesendoderm) related cells. In contrast, the EME element was more accessible in paraxial mesoderm 1, cardiac mesoderm, and mesenchyme cells (Fig. 1h). Thus, while both Neo_ME and EME elements regulate *Mesp1* expression, there may exhibit distinct usage preferences of regulatory element among distinct celltypes.

To further explore the role of the newly-identified element, we performed enhancer reporter assays, demonstrating that the specific distribution of the Neo-ME element was consistent with *Mesp1* expression (Extended Data Fig. 3g). Additionally, we established ESC cell lines with specific deletion of the Neo-ME element and EME element, respectively (Extended Data Fig. 3h). As expected, genetic removal of these two elements led to specific downregulation of *Mesp1* expression during embryoid bodies (EBs) differentiation (Extended Data Fig. 3h). Thus, the newly-identified Neo_ME element is likely a critical regulatory element responsible for *Mesp1* expression.

Together, through single-cell co-profiling of gene expression and chromatin accessibility in the mouse gastrula, combined with the gene-peak linkage capturing strategy (BioCRE), we established a comprehensive multi-omic atlas which captures the celltype-specific GRNs encompassing gene expression, chromatin accessibility of regulatory elements, as well as gene-peak linkage underlying mouse gastrulation.

### Spatiotemporal multi-omic landscape of the *in-silico* gastrulating cells

Recent technological advances have enabled the measurement of gene expression across tissue sections using various spatial transcriptomic strategies.^9–13^ However, most studies covered only the transcriptomic data module and provided limited real 3D spatial context. A comprehensive spatiotemporal multi-omic map that contain genuine spatiotemporal coordinates for embryo tissues remains elusive.

To construct a spatiotemporal multi-omic map that incorporates the spatial coordinates and time stamps of the cells of the gastrulating mouse embryo, we leveraged the published spatiotemporal registered transcriptome dataset of mouse embryonic tissues during gastrulation^5,7^ and the spatial transcriptome reconstruction algorithm-Tangram.^14^ We refined a pipeline (see *Methods*) to reconstruct and visualize celltype distribution, single-cell based spatial transcriptome, and spatiotemporally resolved chromatin accessibility information from embryonic cells at the registered spatiotemporal resolution (Fig. 2a; Extended Data Fig. 4a) in a ST-MAGIC atlas (SpatioTemporal reconstructed Multi-omic Atlas of the Gastrulating In-silico Cells). Beyond spatial distribution, ST-MAGIC can be customized to investigate the spatial distribution of specific celltypes, gene expression, and chromatin accessible domains (Fig. 2a). It is noteworthy that different from the previous gene activity score based chromatin accessibility spatial mapping,^4^ the integrative usage of common cell barcodes present in both the transcriptome and chromatin accessibility modules should significantly enhance the spatial accuracy of the ST-MAGIC.

**Fig. 2.**
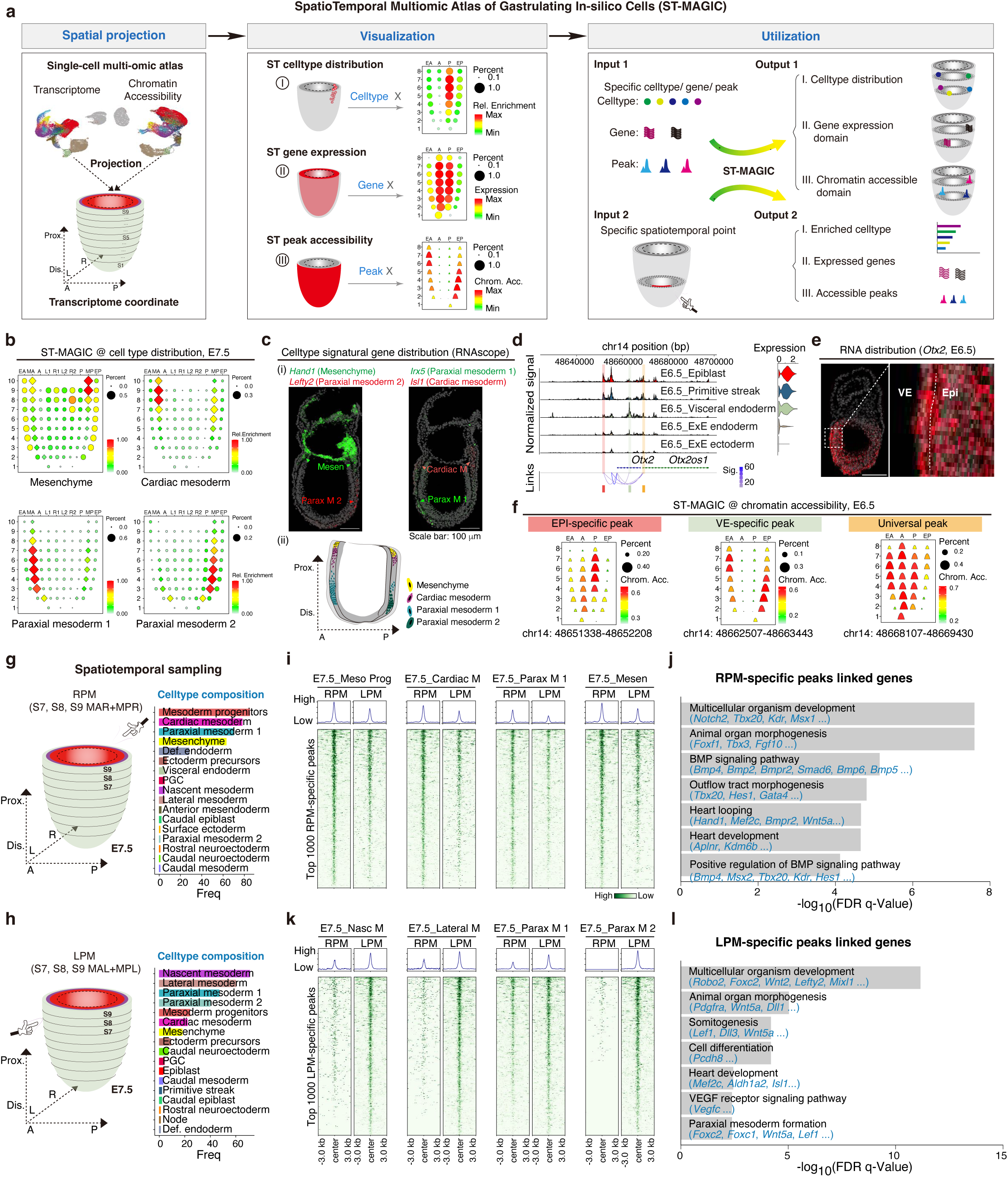
Spatiotemporal tracing of the cellular, transcriptomic, and epigenetic features of cells in the gastrulating embryo by ST-MAGIC. **a.** Schematic of the ST-MAGIC, for spatiotemporal projection and reconstruction of multi-omic atlas at the stage-matched spatial coordinates defined by transcriptome^5,7^. In the reconstructed map (corn-plots), registered positions are labelled in the epiblast/ectoderm domain (A, anterior; L, left lateral; R, right lateral; L1/R1, left/right anterior lateral, L2/R2, left/right posterior lateral), the prospective primitive streak domain (P), the prospective mesoderm domain (M, mesoderm; MA, anterior mesoderm; MP, posterior mesoderm), and the prospective endoderm domain (E, endoderm, EA, anterior endoderm; EP, posterior endoderm) were labelled at the top of each corn-plot. Numbers representing the positions along with the proximal-distal axis in a descending series were labelled on the left side of each corn-plot. Percent represents the ratio of searched celltype, the ratio of expressed genes or the ratio of opened peaks in cells mapped to indicated spot. The relative abundances of enriched celltype, gene expression level, or chromatin accessibility level were shown on the right side of each corn-plot. Rel. enrichment is short for relative enrichment, Chrom. Acc. is short for chromatin accessibility level. To visualize the multi-omic information, spots in corn-plot for celltype and gene expression reconstruction were outlined as round circles, while spots in corn-plot for chromatin accessibility reconstruction were outlined as peak shapes. For sourcing information from the so-constructed ST- MAGIC atlas, we enables data query with a specific cell-type, gene/peak input or starts from specific spatiotemporal points. **b.** The corn-plot displaying the spatial distribution of selected mesodermal celltypes as indicated by the ST-MAGIC atlas at the E7.5 mouse embryo. **c. (i)** The spatial distribution of specific mesodermal cell types validated by expression of marker genes by RNAscope analysis. Mesen: mesenchyme; Parax M 1: paraxial mesoderm 1; Parax M 2: paraxial mesoderm 2; Cardiac M: cardiac mesoderm. (**ii**) Schematic summarizing the distribution of mesoderm subtypes. **d-f.** ST-MAGIC identifying spatial-specific usage of regulatory elements for widely expressed genes. **d.** Representative genome browser snapshot showing the chromatin accessibility around the *Otx2* locus for E6.5 cells. The expression of *Otx2* was also shown on the right. **e.** RNAscope validating the localization of *Otx2* transcripts in E6.5 embryos. **f.** ST-MAGIC visualization of the EPI-specific, VE-specific peak and universal peak linked to *Otx2* expression. VE is short for visceral endoderm, Epi is short for epiblast. **g. h.** Region-specific exploration of the ST-MAGIC atlas of E7.5 mouse embryo identifying the multi-omic basis of left-right symmetry breaking. To capture differences between left and right mesoderm tissues, the left-right resolved GEO-seq transcriptomic coordinate reference was used ^5^. Precise digital spatial sampling of the right (**g**) and left (**h**) proximal mesoderm (RPM and LPM) captures the lateral region celltype composition in the E7.5 mouse embryo. **i. j.** Heatmaps showing the chromatin accessibility of top 1000 specific accessible peaks derived from the RPM across the major celltypes enriched in the RPM region (**i**). The chromatin accessibility profiles of the same peakset from the celltype counterparts of the counter-lateral (LPM here) were also shown. Gene ontology analyses of RPM accessible peaks-linked genes were also performed (**j**). Meso Prog is short for mesoderm progenitors; Cardiac M is short for cardiac mesoderm; Parax M 1 is short for paraxial mesoderm 1; Mesen is short for mesenchyme. **k. l.** Heatmaps showing the chromatin accessibility of top 1000 specific accessible peaks derived from the LPM across the major celltypes enriched in the LPM region (**k**). The chromatin accessibility profiles of the same peakset from the celltype counterparts of the counter-lateral (RPM here) were also shown. Gene ontology analyses of LPM accessible peaks-linked genes were shown (**l**). Nasc M is short for nascent mesoderm; Lateral M is short for lateral mesoderm; Parax M 1 is short for paraxial mesoderm 1; Parax M 2 is short for paraxial mesoderm 2.

To assess the efficacy of the ST-MAGIC, JSD indexing revealed that a high consistency of the genome-wide transcriptomic architecture gleaned from the ST-MAGIC with that from the GEO-seq dataset (Extended Data Fig. 4b), and cells were mapped evenly across spatial spots in the constructed atlas (Extended Data Fig. 4c). Besides, taking the expression of *Sox2* as an example, the expression pattern of *Sox2*, gleaned from the ST- MAGIC, closely matched the GEO-seq resource and also the experimental results (Extended Data Fig. 4d, 4e). Histone modification H3K27ac has been usually used as an active histone marker frequently deposited at accessible chromatin regions.^15^ Following this, we selected chromatin regions with region-specific H3K27ac distribution, as revealed in our previous study,^3^ and checked their spatial distribution of chromatin accessibility from the ST-MAGIC. For example, the chromatin region (chr10: 63347793-63348989), which locates upstream of the *Sirt1* gene and is specially marked by H3K27ac in the anterior epiblast (A) region (Extended Data Fig. 4f), also exhibits anterior epiblast-specific accessible pattern as revealed by the ST-MAGIC (Extended Data Fig. 4f).

We further analyzed the spatial distributions of the embryonic celltypes (Extended Data Fig. 5a). Mostly, celltypes were assigned to their expected embryonic spatial positions (Extended Data Fig. 5a-c). For example, ectoderm precursors were mapped to the anterior region of the epiblast, while primitive streak cells were specially mapped to the expected posterior region (Extended Data Fig. 5a). Interestingly, we observed a clear regionalization of the mesoderm subtypes, particularly in the E7.5 embryos (Fig. 2b; Extended Data Fig. 5b, 5c). Specifically, the cardiac mesoderm and mesenchyme cells were mostly located in the proximal region, while the paraxial mesoderm 1 cells and the paraxial mesoderm 2 cells were situated in the medial-distal region (Fig. 2b). The spatial distributions for these mesoderm subtypes were validated by determining the related signatural gene expression *in vivo* (Fig. 2c; Extended Data Fig. 5d, 5e).

Apart from revealing the formation of celltype-specific spatial territories, ST-MAGIC also unveils the spatial-specific usage of distal linked peaks for gene regulation. For example, *Otx2*, which is broadly expressed in the epiblast (EPI) and visceral endoderm (VE) ^16^ (Fig. 2d, 2e), is linked to three spatial types of chromatin peaks: EPI-specific, VE-specific, and universal peaks (Fig. 2f; Extended Data Fig. 5f). To specify, ST-MAGIC revealed that EPI-specific peaks were predominantly located in the epiblast region, while the VE-specific peaks were located in the visceral endoderm region (Fig. 2f; Extended Data Fig. 5f).

ST-MAGIC allows the exploration of specific celltype compositions, gene expression pattern, and chromatin accessibility at any spatial or temporal coordinate across mouse gastrulation (Fig. 2a). Previously, we reported the symmetry breaking event for the left-right body axis first emerges at the late gastrulation stage, manifesting as differential BMP signaling activity and target gene expression in the contralateral proximal mesoderm.^5^ To trace the multi-omic basis for the initiation of left-right asymmetry, we extracted the molecular information from the proximal lateral region of the mesoderm layer using ST- MAGIC (Fig. 2g-l). Consistently, we found that genes related to left-right symmetry breaking also showed laterally biased expression (Extended Data Fig. 6a). Exploration of celltype composition revealed that the mesoderm progenitors, cardiac mesoderm, paraxial mesoderm 1 and mesenchyme cells were over-represented in the right proximal mesoderm region (RPM), while nascent mesoderm, lateral mesoderm, paraxial mesoderm 1 and paraxial mesoderm 2 cells were enriched in the left proximal mesoderm region (LPM) (Fig. 2g, 2h). Profiling of the top 1000 specific accessible peaks in both RPM and LPM regions revealed that asymmetric levels of chromatin accessibility was discernable between left and right celltype counterparts (Fig. 2i, 2k; Supplementary Table 2). To identify the potential biological functions of these asymmetric peaks, we performed gene ontology analyses for these peaks linked genes. Interestingly, BMP signaling pathway-related genes were associated with RPM enriched peaks, while genes related to somitogenesis and heart development were regulated by peaks with higher accessibility in the LPM (Fig. 2j, l). We also traced the emergence of these asymmetric peaks during gastrulation (Extended Data Fig. 6b-j). Intriguingly, both left and right lateral asymmetric peaks became accessible from E6.75 onward (Extended Data Fig. 6b, 6h), coinciding with the emergence of mesoderm subtypes from the primitive streak cells. For example, two specific peaks, linked with *Lefty2* expression became accessible at the E6.75 nascent mesoderm cells when *Lefty2* expression begins, and showed LPM higher distribution at the E7.5 stage when *Lefty*2 expression is higher on the left side (Extended Data Fig. 6c-f). One of the two peaks is related to a known ASE element,^17^ while the other is a newly-identified *Lefty2* regulatory element (Neo_LRE) (Extended Data Fig. 6e). Genetic deletion of Neo_LRE in mouse embryonic stem cells showed that *Lefty2* expression was severely affected in the knockout (KO) cells compared to wild-type (WT) controls during gastruloid differentiation *in vitro* (Extended Data Fig. 6g).

Thus, the ST-MAGIC resource established by spatiotemporal reconstruction of the multi-omic atlas to temporal-matched spatial coordinates facilitates in-depth investigation of multi-dimensional molecular architectures of celltypes in defined domains of the gastrulating embryo.

### ST-MAGIC (+) infers the spatiotemporal turn-over of enhancer regulons

The hierarchical activation of gene regulatory networks by developmental signals and TFs is crucial for mammalian development.^18^ To explore the logics of TF GRNs underpinning gastrulation, we applied the SCENIC+,^19^ a method for the inference of enhancer-driven TF GRNs, to systematically profile the TF distribution, TF target gene expression as well as TF target peak chromatin accessibility across the annotated celltypes. Major regulators for germ layer development were recovered (Fig. 3a; Extended Data Fig. 7a). Notably, while the distribution of TF expression showed high celltype-specificity, the presence of TF targets especially for the TF target peaks, often exhibited shared pattern among closely related neighbors in the same pedigree (Fig. 1c, 3a). For example, in the blood cell lineage, which encompasses hematoendothelial progenitors, blood progenitors 1 and blood progenitors 2, a clear TF expression hierarchy from *Etv2*, *Gata2* to *Tal1* was observed. However, the chromatin accessibility for these TF targets remain indistinguishable. This phenomenon supports that *Etv2* could function as a priming factor responsible for enhancer opening prior to the hematoendothelial fate commitment (Fig. 3a).^20^ The sharing of accessible TF target peaks (such as *Noto* TF) was also detected in cells of axial mesendoderm lineage and definitive endoderm lineage (Fig. 3a), which were both differentiated from the anterior primitive streak (Fig. 1c).

**Fig. 3.**
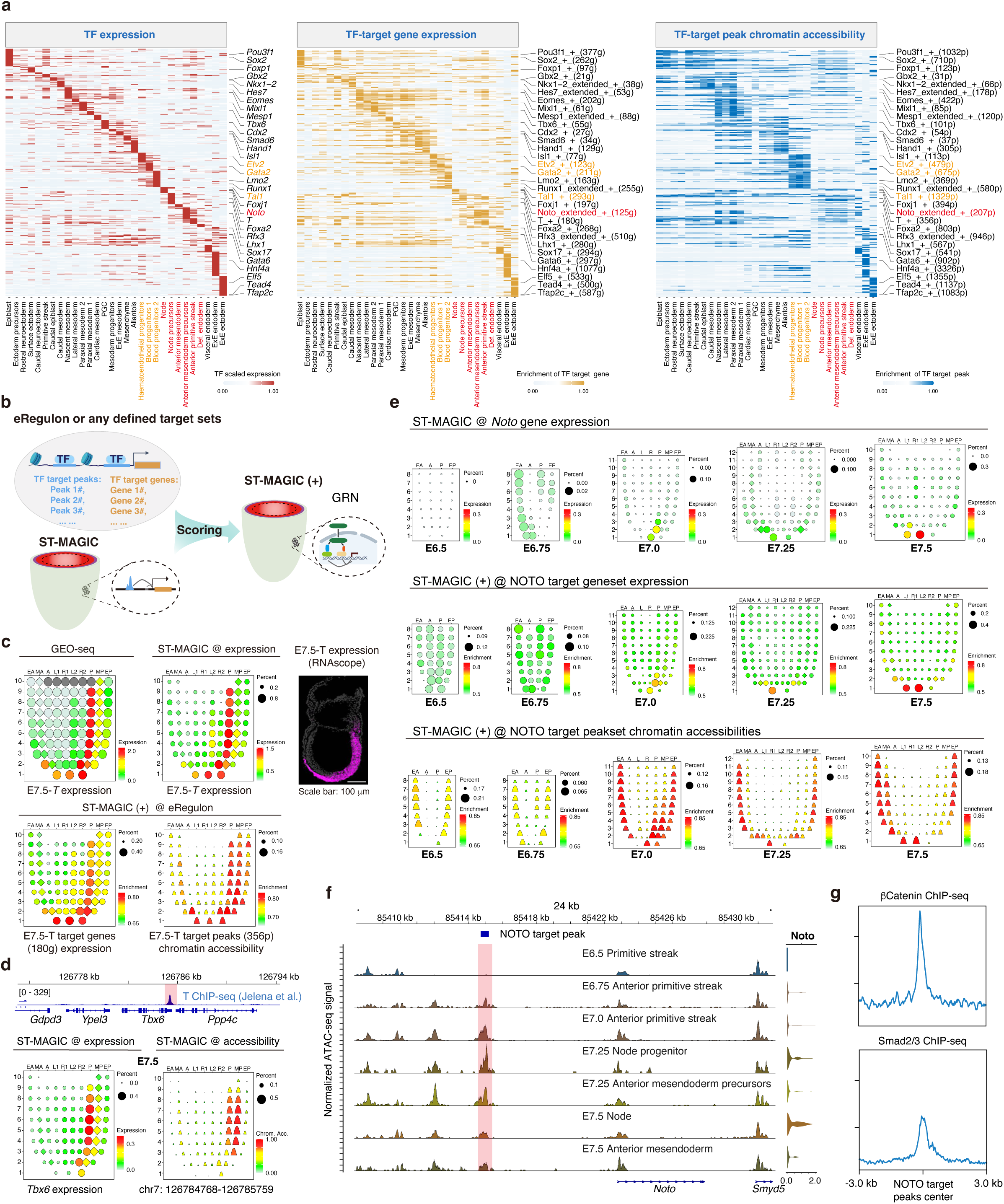
Expanded ST-MAGIC enables inference of eRegulon dynamics during gastrulation. **a.** Heatmap showing the transcription factor (TF) expression (left), the AUC score of TF- target geneset (middle), and the AUC score of TF-target peakset (right) of the eRegulon on a color scale. Celltypes were ordered on the basis of their developmental relationship. Representative TF eRegulons were labelled next to the heatmap. **b.** Scheme of the workflow to generate the expanded version of ST-MAGIC. **c.** The visualization of *T* expression and the downstream regulatory network in E7.5 embryo using ST-MAGIC and ST-MAGIC (+). The distribution of *T* transcripts was validated by RNAscope (right panel). **d.** Representative ST-MAGIC snapshot of T targets in E7.5 embryo. *Tbx6* and its genomic neighboring peak bound by T were specifically selected. T ChIP-seq data was drawn from the published dataset.^50^ **e.** ST-MAGIC (+) visualization of NOTO eRegulon revealing the spatiotemporal asynchrony among *Noto* expression, NOTO target gene expression and NOTO target peak chromatin accessibility. **f.** Genome browser snapshot displaying the dynamics of chromatin accessibility of NOTO target peak (blue square marked region) around *Noto* locus with corresponding gene expression level (right side). **g.** Enrichment of β-Catenin (top) and Smad2/3 (bottom) at NOTO target chromatin peaks. ChIP-seq information was drawn from dataset of βCatenin ChIP-seq in EpiLCs^26^ and Smad2/3 ChIP-seq from EBs^29^.

To explore the spatiotemporal distribution of these eRegulon imputed by SCENIC+, we developed a pipeline called ST-MAGIC (+), which projects the pre-defined TF target gene sets or target peak sets to the ST-MAGIC atlas through Area Under Curve (AUC) scoring (Fig. 3b). Theoretically, the so-constructed ST-MAGIC (+) should be able to visualize the spatiotemporal turnover of TF GRNs during mouse gastrulation. To validate the fidelity of ST-MAGIC (+) profiling, we first checked the spatial distribution of region-specific H3K27ac modified peaks inferred from the epigenetic landscape of the mouse gastrula.^3^ As shown, region-specific peak sets were accurately mapped to their sampling origins (Extended Data Fig. 8a). Next, we examined the spatial distribution of GRNs for the major regulators identified in Fig. 3a. Taken transcription factor, *T*, as an example, GEO-seq and ST-MAGIC revealed that *T* expression is localized to the primitive streak and anterior mesendoderm in E7.5 embryo (Fig. 3c), and T’s target genes showed a similar distribution pattern (Fig. 3c). However, for the target peaks of T, we observed a broader spatial distribution pattern than the TF expression, manifesting as high level of chromatin accessibility in both the primitive streak and neighboring mesodermal regions (Fig. 3c). This observation was verified through determining the direct T binding peak around the gene locus of *Tbx6*, which is a well-known T target gene, using ST-MAGIC (Fig. 3d). Moreover, considering that T continues to express in the primitive streak throughout gastrulation and that the mesoderm cells are the immediate progeny of the primitive streak cells,^21^ the broad accessible pattern of T binding chromatin regions suggests that T may act as a priming factor with the ability to open a broad spectrum of chromatin regions and instruct the subsequent fate commitment of mesoderm cells.

Finally, we dissected the dynamics of shared accessible patterns among trajectory neighborhoods for the axial mesendoderm lineage. As resolved by ST-MAGIC (+), we found that the target genes of NOTO exhibit a matched spatiotemporal distribution with the TF expression, but the chromatin accessibilities of NOTO target peaks were widespread in the endoderm cell layer of the E7.5 embryo, where *Noto* is not expressed (Fig. 3e). Further examination revealed that the chromatin accessibility of NOTO target peaks was elevated at early gastrulation stage, especially in the E7.0 anterior primitive streak region, well before *Noto* expression emerged (Fig. 3e; Extended Data Fig. 8b). Detailed exploration of the direct NOTO target chromatin region around the *Noto* locus supported the spatiotemporal kinetics of NOTO eRegulon by ST-MAGIC (+) (Fig. 3f). This result indicates that crucial upstream players would be involved in setting up the chromatin level of NOTO GRN during axial mesendoderm lineage development. It has been reported that the modulations of WNT and NODAL signaling are involved in the anterior primitive streak patterning.^22^ We also found that both WNT and NODAL signaling are highly enriched at these chromatin regions (Fig. 3g), which suggesting crucial inductive signals may participate in establishing the pre-accessible pattern for NOTO targets.

### ST-MAGIC (+) reflects the spatiotemporal responsiveness to developmental signaling

Inductive developmental signals such as WNT and NODAL signaling, act as morphogen cues with localized secretion but distant function, playing crucial roles in elaborating the proper embryo arrangement and related cell lineages. Multi-layered dimensions, including the morphogen concentration, duration, the completeness of the signaling cascade components, and the intrinsic competence of the receiving cells influence interpretation of signals. Currently, a systematic evaluation of the intrinsic chromatin competence for signaling function remain unexplored.^23^ To characterize the spatial context-dependent mechanisms for developmental signals interpretation, we applied ST-MAGIC (+) to investigate the hierarchy of signal and effector distribution, associated chromatin responsiveness and gene-peak linkage inferred transcription output along the lineage trajectory (Fig. 4a). Generally, the spatial distributions of key signal components were characterized by determining the expression pattern of related genes, the chromatin responsiveness to signals was delineated by profiling the spatial accessibility of direct signal effector targets, and the spatial transcription output were measured by checking the expression pattern of signal effector chromatin binding regions linked genes (Fig. 4a).

**Fig. 4.**
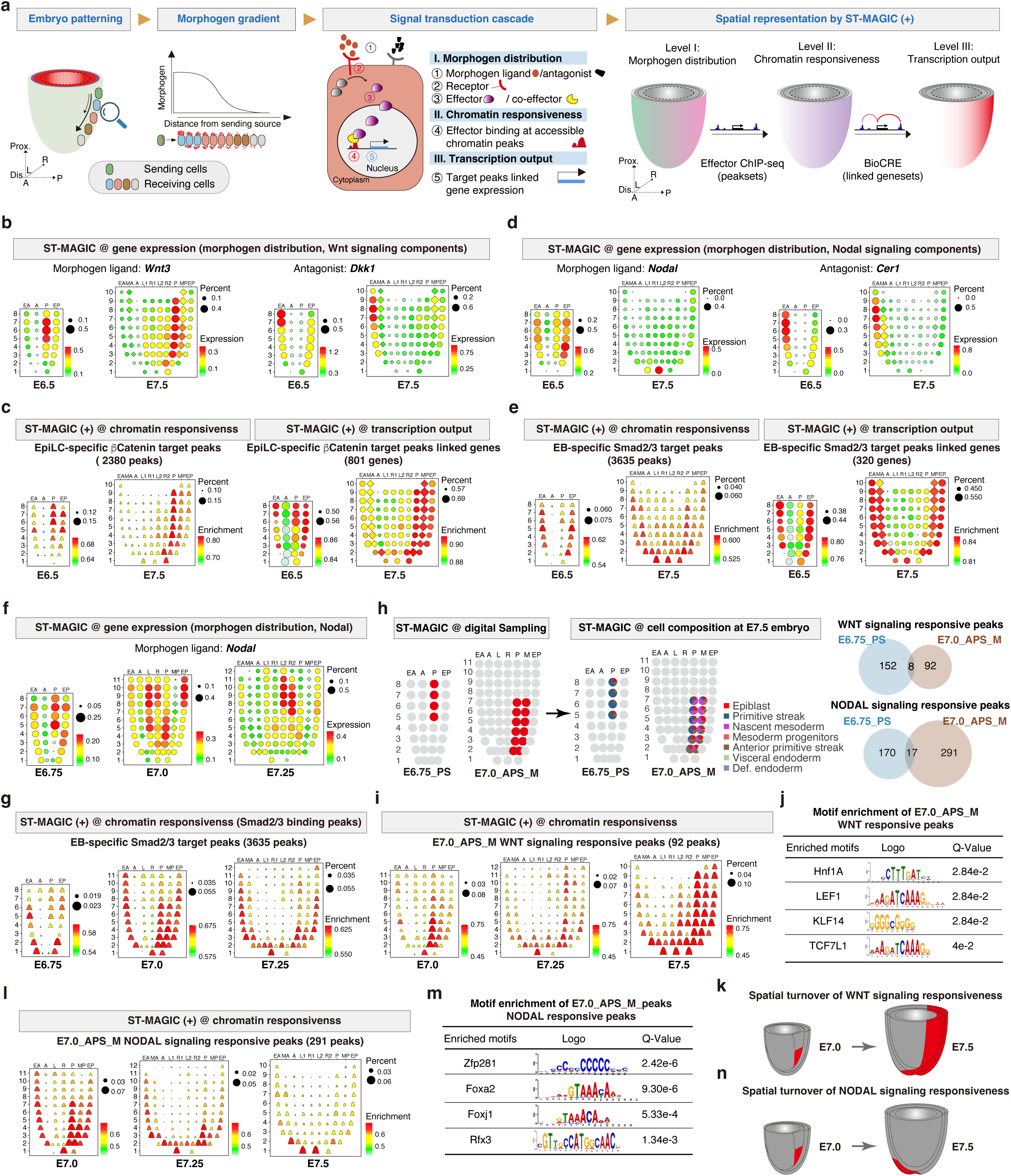
ST-MAGIC (+) demarcates the spatiotemporal changes in responsiveness to signaling pathways. **a.** Schematic overview of the morphogen gradient and the interpretation of signal transduction cascade in embryo patterning, and the *in silico* visualization using ST-MAGIC (+). **b.** ST-MAGIC visualization of the distribution of WNT signaling ligand-*Wnt3* (left) and the antagonist-*Dkk1* (right) during mouse gastrulation. **c.** The 2380 EpiLC-specific βCatenin binding peaks and the 801 linked genes were mapped to E6.5 and E7.5 embryo to show the spatiotemporal dynamics of chromatin responsiveness (left) and transcriptional output of WNT signaling (right). **d.** ST-MAGIC visualization of the NODAL signaling ligand-*Nodal* and the antagonist-*Cer1* during mouse gastrulation. **e.** ST-MAGIC (+) visualization of embryoid body (EB)-specific Smad2/3 binding peaks (3635 peaks, left) and the linked genes (320 genes, right) reflecting the spatiotemporal transition of intrinsic chromatin responsiveness to NODAL signaling. **f. g.** The spatiotemporal expression of *Nodal* (**f**) and the transition of intrinsic chromatin responsiveness to NODAL signaling (**g)**. **h.** The transition of intrinsic chromatin responsiveness to signaling is mostly related to E6.75_PS region (enriched with primitive streak cells) and E7.0_APS_M region (enriched with anterior primitive streak and nascent mesoderm). Left: the specific spatial locations of E6.75_PS and E7.0_APS_M region; middle: the celltype composition at indicated regions; top right: venn diagram showing the direct comparison of region-specific WNT signaling responsive peaks between E6.75_PS and E7.0_APS_M; bottom right: venn diagram showing the direct comparison of region-specific NODAL signaling responsive peaks between E6.75_PS and E7.0_APS_M. **i. j.** ST-MAGIC (+) visualization (**i**) and motif enrichment (**j**) of the 92 E7.0 APS_M specific WNT signaling responsive peaks during the late stage (E7.0, E7.25, E7.5) of mouse gastrulation. **k.** The model describing the spatial evolution of WNT signaling responsiveness during the late stage of mouse gastrulation. **l. m.** ST-MAGIC (+) visualization (**l**) and motif enrichment (**m**) of the 291 E7.0 APS_M specific NODAL signaling responsive peaks during the late stage (E7.0, E7.25, E7.5) of mouse gastrulation. **n.** The model describing the spatial evolution of NODAL signaling responsiveness during the late stage of mouse gastrulation.

We first analyzed the global pattern and the cellular responsiveness of WNT signaling across gastrulation (Extended Data Fig. 9a). Consistent with known biology,^24^ the Wnt signal ligand-*Wnt3* was predominantly expressed in the primitive streak and adjacent mesoderm tissue, while the Wnt signal antagonist-*Dkk1* was specifically expressed in the anterior visceral endoderm region (Fig. 4b). However, the WNT signaling receptor-*Lrp6* and the effector-*Ctnnb1* (also known as β-Catenin) were widely distributed within the gastrula (Extended Data Fig. 9b).

To systematically characterize the embryo chromatin responsiveness to WNT signaling, we incorporated two published ChIP-seq datasets for WNT signaling effector-β-Catenin from the *in vitro* ESCs^25^ and EpiLC^26^. Through differential binding peaks analyses, the chromatin regions were classified into three groups of common, ESC-specific, and EpiLC-specific peaks (Extended Data Fig. 9c, 9d). Spatiotemporal reconstruction of these peaks though ST-MAGIC (+) revealed that ESC-specific peaks were accessible in the anterior epiblast, where cells remain in a pluripotent state^27^ (Extended Data Fig. 9e). Meanwhile, for the peaks with EpiLC-specific β-Catenin binding, ST-MAGIC (+) reported that these peaks were accessible in the embryonic region largely overlapped with *Wnt3* distribution (Fig. 4b, 4c). In-depth analyses of the ST-MAGIC+ results for EpiLC-specific peaks across five stages of the gastrula showed that the PS region and neighboring posterior mesoderm region remained consistently accessible across the gastrulation (Fig. 4b, 4c; Extended Data Fig. 9f). Together, these results underscore that in response to WNT signaling input, cells first turnover the Wnt-responsive chromatin landscape during the transition from pluripotent state to lineage-specified progenitors, and then modulate the chromatin accessibility to accommodate the posterior regionalized mesoderm subtypes during gastrulation (Extended Data Fig. 9g).

We next analyzed the intrinsic cellular responsiveness to NODAL signaling (Extended Data Fig. 10a), a crucial signal for embryo patterning and germ layer formation.^28^ Examination of the NODAL signaling components in the ST-MAGIC atlas revealed that the ligand-Nodal was initially expressed in the posterior epiblast at early gastrulation stage but abruptly shifted to the distal tip region at the E7.5 stage (Fig. 4d). In the meantime, the NODAL signaling antagonist-*Cer1* was constantly expressed in the anterior visceral endoderm region (Fig. 4d, Extended Data Fig. 10c). However, the NODAL signaling receptors and the effectors were widely expressed in the gastrula (Extended Data Fig. 10b). The changes of expression domain of the NODAL ligand-antagonist pair should lead to the corresponding shift of the extrinsic NODAL morphogen gradient during mouse gastrulation.

To chart the chromatin responsiveness along with the NODAL pattern shift, we incorporated the published ChIP-seq datasets for NODAL signaling effectors SMAD2 and SMAD3.^29^ We found that the chromatin binding ability of Smad2/3 complex was largely *de novo* generated in EB cells, an *in vitro* counterpart of mesoderm cells (Extended Data Fig. 10d, 10e). GREAT analyses further indicated that these peaks are involved in the cell fate specification, anterior/posterior formation and primitive streak formation (Extended Data Fig. 10e). Then, we used ST-MAGIC (+) to profile the spatial distribution of these peaks during gastrulation. Interestingly, we found that the global chromatin accessibilities of these EB-specific Smad2/3 binding peaks first appeared in both the endoderm region and PS regions (Fig. 4e). Subsequently, the accessibility of these peaks in the PS region gradually shifted and formed a distal-to-proximal accessibility gradient in the E7.5 gastrula (Fig. 4e; Extended Data Fig. 10f). Notably, the chromatin responsiveness to Nodal signaling began relocating at an earlier stage between E6.75 and E7.0, ahead of the re-alignment of the Nodal gradient in the germ layers occurring between E7.25 and E7.5 stage (Fig. 4f, 4g). These results highlight that the spatial turnover of NODAL signaling chromatin responsiveness occurs earlier than the re-positioning of *Nodal* activity.

To determine the intrinsic features for the relocation of signaling chromatin responsiveness, we extracted the specific subset of signaling responsive chromatin peaks which show high accessibility at the E6.75 primitive streak region (E6.75_PS) or at the E7.0 anterior primitive streak and adjacent mesoderm region (E7.0_APS_M) (Fig. 4h). Generally, the signaling responsive peaks for NODAL and WNT signaling were largely independent between E6.75_PS region and E7.0_APS_M region (Fig. 4h; Supplementary Tables 3, 4). As shown, the E7.0_APS_M specific WNT signaling responsive peaks were gradually extended to the whole posterior region (Fig. 4i), and enriched with typical WNT signaling co-effector LEF1 and TCF7L1 motifs (Fig. 4j). The posterior enriched WNT signaling responsiveness (Fig. 4k) support the notion that WNT signaling is instrumental for regulating posterior embryo development.^23^ Interestingly, for NODAL signaling, profiling of the E7.0_APS_M specific signal responsive peaks exhibited a gradual shift pattern of these peaks towards the distal tip at E7.5 embryo, where *Noto* and *Nodal*-expressing cells reside (Fig. 3e, 4l). Functional characterization of E7.0_APS_M NODAL signaling responsive peaks showed enrichment in chordate embryonic development and embryo pattern specification process, and the knockout of related genes can frequently lead to abnormal germ layer morphology, abnormal rostral-caudal axis patterning and absent floor plate (Extended Data Fig. 10g). Motif enrichment analyses of these peaks revealed significant enrichment of major TF regulators especially for axial mesendoderm development such as *Zfp281*, *Foxa2*, and *Foxj1* (Fig. 4m). These results indicate that the spatial relocation of embryonic responsiveness to Nodal signaling may play specific roles in the forthcoming cell development of axial mesendoderm lineage (Fig. 4n).

### Molecular hierarchy underlying axial mesendoderm lineage development

The axial mesendoderm has been reported to be the direct developmental antecedent for the midline notochord cells, which instructs the following somitogenesis and neural patterning.^30^ To better understand axial mesendoderm cell development, it is essential to document the spatiotemporal context of stepwise appearance of molecular features and cellular states during the lineage formation process, with accurate spatiotemporal information.

Here, based on the inferred lineage trajectory for the gastrula (Fig. 1c, 5a) and the ST- MAGIC reconstructed cell distribution for the mouse gastrula (Extended Data Fig. 5a), we found that cells residing in the E7.5 distal tip, where the prospective notochord cells located,^31^ are composed of the node cells and anterior mesendoderm cells (Fig. 5a; Extended Data Fig. 11a). Molecular identification of known notochord markers, *Shh*, *Noto*, and *Foxj1*, revealed that the presence of two distinct cell subtypes (*Shh*^+^*Noto*^-^*Foxj1*^-^ and *Shh*^+^*Noto*^+^*Foxj1*^+^) at the ventral distal surface for the E7.5 embryo, and the two distinct cell subtypes can persist through organogenesis (Fig. 5b; Extended Data Fig. 11b, c).

**Fig. 5.**
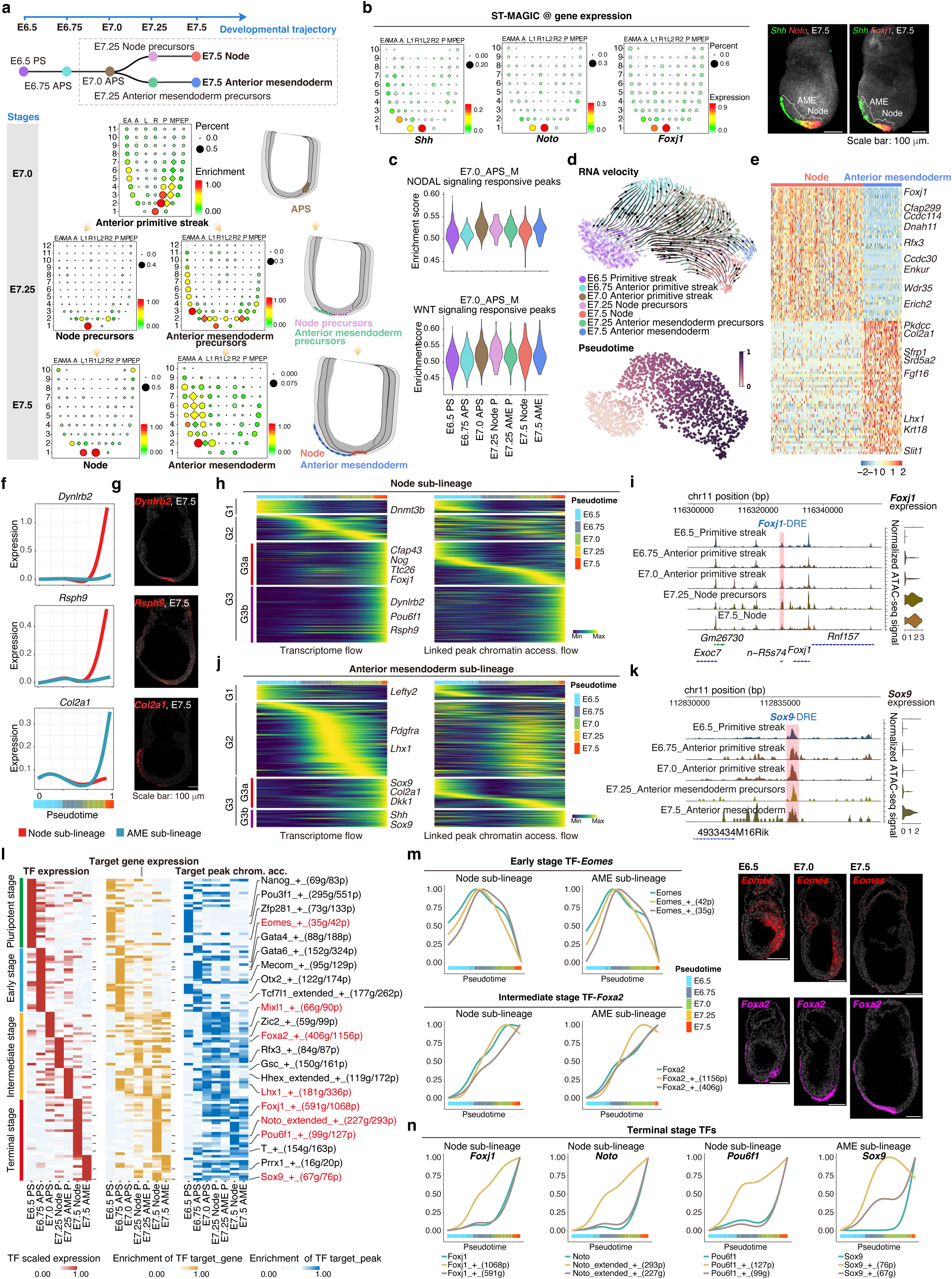
The sequential process of cell fate commitment in the axial mesendoderm lineages. **a.** ST-MAGIC revealing the spatiotemporal information for lineage trajectory of axial mesendoderm (top) from the anterior primitive streak cells of E7.0 embryo to the derivatives – the node and anterior mesendoderm of E7.5 embryo.. Bottom: ST-MAGIC results showing the changes in the location of axial mesendoderm lineage-related cells during gastrulation. PS is short for primitive streak cells, APS is short for anterior primitive streak cells. **b.** ST-MAGIC showing the expression of known markers for node and anterior mesendoderm cells (*Shh*, *Noto*, *Foxj1*), which were validated by whole-mount RNAscope. AME is short for anterior mesendoderm. **c.** Violin plot showing the AUC scoring for E7.0_APS_M NODAL responsive peaks (top) and WNT signaling responsive peaks (bottom) accessibilities during axial mesendoderm development. Node P is short for node precursors, AME P is short for anterior mesendoderm precursors. **d.** UMAP of mouse axial mesendoderm during gastrulation with scVelo-projected velocities, shown as streamlines. Top: UMAP representation with RNA velocity stream plots overlaid for axial mesendoderm lineage; bottom: UMAP representation with inferred trajectories colored by pseudo-time ordering. **e.** Heatmap showing the direct comparison of differentially expressed genes for the two axial mesendoderm subtypes (node and anterior mesendoderm). **f. g.** Line chart describing the smoothened gene expression trends in pseudotime (**f**). The trend for each gene was shown for each trajectory leading to the terminal populations of node (*Dynlrb2* and *Rsph9*) or anterior mesendoderm (*Col2a1*). The gene expression pattern was validated by RNAscope (**g**). **h-k.** Heatmaps displaying the smoothened gene expression trends (left) and chromatin accessibility levels of linked peaks (right) in node (**h. i.**) and anterior mesendoderm development (**j. k.)**. Cells ordered by the pseudo-time were marked (top of each heatmap). **l.** SCENIC+ analyses revealing the transcription factor hierarchy during axial mesendoderm development. **m.** Line plot describing the TF expression, TF-target gene expression, as well as TF-target peak accessibility of Eomes (top) and Foxa2 (down) during axial mesendoderm lineage. The spatiotemporal expression dynamics for both TF was validated at the right. **n.** Line plot describing the TF expression, TF-target gene expression, as well as TF-target peak accessibility for *Foxj1*, *Noto*, *Pou6f1*, and *Sox9* during axial mesendoderm development.

The inferred lineage trajectory indicates that the node and the anterior mesendoderm cells are spatially specified from the E7.0 APS cells (Fig. 5a). Concurrently, chromatin responsiveness to NODAL and WNT signaling gradually increased in the E7.0 anterior primitive streak cells and is maintained at high accessibility levels till the E7.5 stage (Fig. 5c). To chart the molecular dynamics along the developmental trajectory, we applied CellRank^32^ to assign the cell fate probabilities for all the related cells at single-cell resolution. RNA velocity and pseudotime ordering revealed that the cells at an early stage (E6.5 primitive streak, E6.75 anterior primitive streak) are unspecified for cell fate directions, while cells at more advanced stages show directed flow towards the E7.5 node and E7.5 anterior mesendoderm (Fig. 5d). Transcriptomic comparison of the two cell subtypes showed that the node cells express high levels of cilia and dynein-related genes, such as *Cfap299* and *Dnah11*, while the anterior mesendoderm cells express a high level of collagen-related genes, such as *Col2a1* (Fig. 5e; Extended Data Fig. 11d). Molecularly, smoothened gene expression trends along the pseudotime order showed that cilium related genes, such as *Dynlrb2* and *Rsph9*, were gradually upregulated down the road to node, while the collagen gene *Col2a1* was expressed till the terminal stage of the anterior mesendoderm (Fig. 5f, g).

Combinatorial pseudotime-based tracking of gene expression cascades and associated chromatin peaks revealed three successive stages (G1, G2, G3) enriched in the early, intermediate and terminal populations, respectively (Fig. 5h-k). Intriguingly, for both lineages, a specific subset of the terminal stage expressed genes (G3a) was regulated by chromatin peaks which get accessible at an earlier stage (Fig. 5h-k). For example, in node cells, *Foxj1* expression was evidently upregulated at the E7.25 stage, but the chromatin accessibility for the linked distal peak-*Foxj1*-DRE was getting accessible in the E7.0 anterior primitive streak cells (Fig. 5i). Similarly, for anterior mesendoderm cells, genes like *Sox9* showed the same pattern (Fig. 5k). These pre-accessible chromatin peaks suggest epigenomic priming for the development of axial mesendoderm lineage.

To systematically determine the molecular hierarchy of the enhancer-driven TF GRNs for the axial mesendoderm lineage, we used SCENIC+ to infer the potential candidate TFs and the dynamics of downstream target genes and linked chromatin peaks (Fig. 5l). A clear hierarchy of TF usage turnover can be captured. To specify, pluripotency-related TFs, such as Nanog, were enriched at the E6.5 stage; early lineage factors, such as *Eomes*, were enriched at the E6.75 stage; subsequently, for the intermediate stage factors, such as *Mixl1, Foxa2* and *Lhx1*, were enriched in the E7.0 anterior primitive streak cells, E7.25 node precursors, and E7.25 anterior mesendoderm precursors, respectively; and terminal stage factors, such as *Noto* and *Sox9*, were abundant in E7.5 node and anterior mesendoderm cells, respectively (Fig. 5l). Detailed analyses of these eRegulons revealed that the early and intermediate TFs first upregulate TFs gene expression, followed by increasing the target gene expression as well as target peak chromatin accessibility (Fig. 5m; Extended Data Fig. 12a). In contrast, for the terminal stage enriched TFs, the chromatin of TF target peaks becomes accessible at an earlier stage, followed by TF expression and TF target gene expression (Fig. 5n). This temporal turnover of TF GRNs may reflect a molecular relay underlying the sequential cascade of cell fate commitment during axial mesendoderm development. Notably, among multiple known TFs GRNs, we identified a novel TF, POU6F1, which exhibits highly specific distribution of both the TF itself and its target genes in the node. However, the chromatin regions targeted by POU6F1 get accessible prior to the expression of the TF and its target genes (Fig. 5n).

### Distinct roles of stage-related TFs in regulating gene expression and setting up chromatin accessibility

To determine the potential significance of TF hierarchy and signal responsiveness during axial mesendoderm lineage development, we first checked the distribution of the TF GRNs for early stage TF (EOMES), intermediate stage TFs (MIXL1, FOXA2, LHX1), and terminal stage TFs (NOTO and POU6F1) during gastrulation using ST-MAGIC (+). Consistent with the SCENIC+ results (Fig. 5m, 5n), we found that for the early and intermediate stage TFs, expression of these TFs commences prior to or concurrently with the target gene expression and target peak accessibility (Extended Data Fig. 12b-e). In contrast, for the terminal stage TFs-NOTO and POU6F1, we observed that their target chromatin peaks first become accessible at the expected APS region and endoderm region at the E7.0 stage, when the TFs and their target genes only show minimal expression (Fig. 4e, 6a, 6b). With the progressing of embryo development, once *Noto* and *Pou6f1* reached the expression summit at the E7.5 node region, the target genes start to show abundant enrichment at the same region (Fig. 4e, 6a, 6b). These results suggest that the early and intermediate stage TFs may function as priming factors by opening a broad spectrum of chromatin regions, thereby setting up a proper chromatin environment which is permissive for the following signaling and the terminal stage TFs-NOTO and POU6F1. The expression of TFs-NOTO and POU6F1 at the terminal stage can then interact with the pre-accessible chromatin regions, and boost the target gene expression and ultimately establish the required transcriptomic state.

**Fig. 6.**
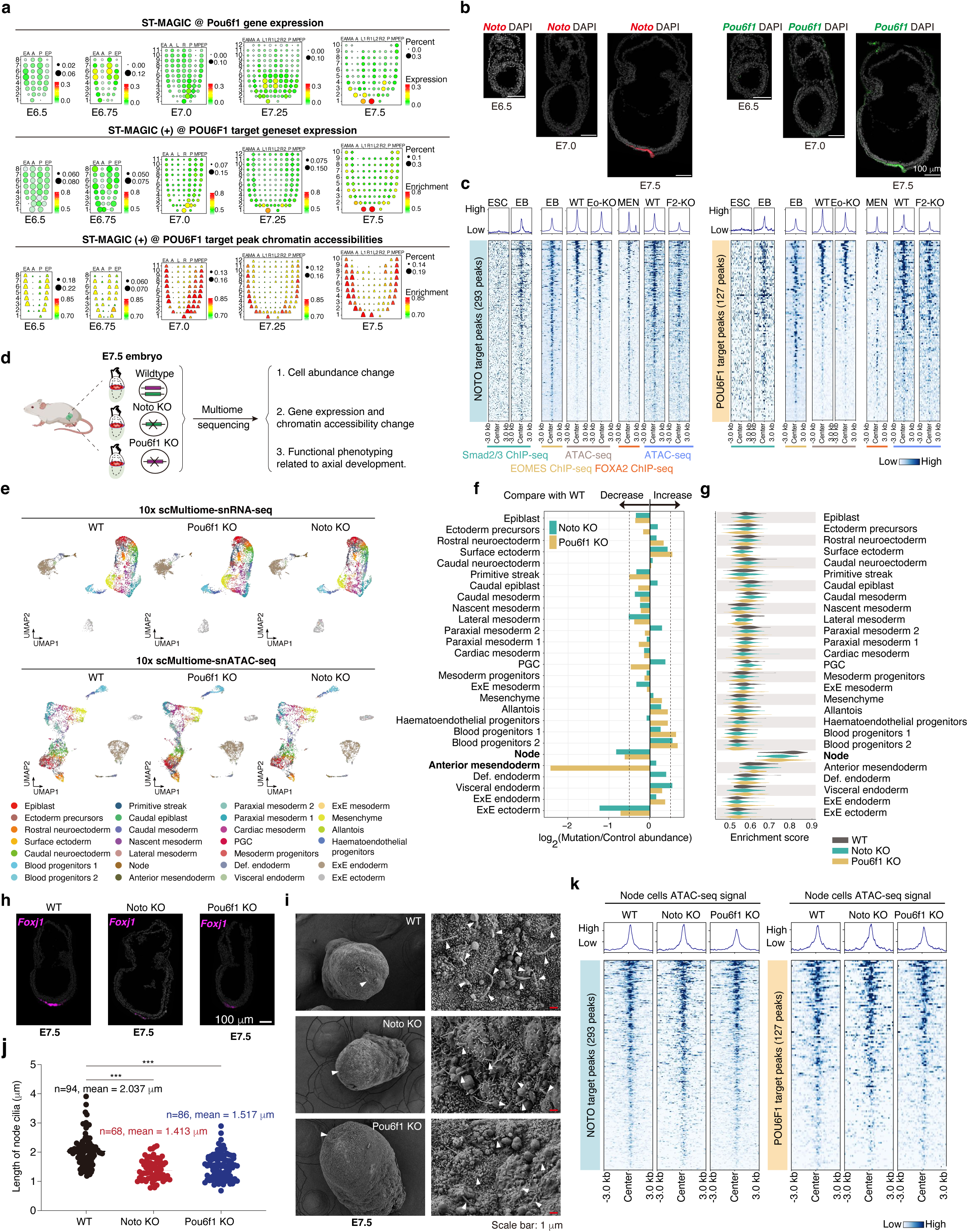
Identification of *Noto* and *Pou6f1* as the last runners of the TF relay during axial mesendoderm lineage development. **a. b.** ST-MAGIC visualizing the distribution of *Pou6f1* eRegulon during mouse gastrulation (**a**). RNAscope experiment was performed to validate the expression of *Noto* and *Pou6f1* in the gastrula (**b**). **c.** Heatmaps showing the enrichment of Smad2/3, EOMES, and FOXA2 binding at the prospective target peak regions of NOTO (left) and POU6F1 (right). Information was sourced from dataset of Smad2/3 ChIP-seq^29^, EOMES ChIP-seq^50^, ATAC-seq for both WT EB cells and Eomes-KO EB cells^51^, FOXA2 ChIP-seq^52^, ATAC-seq for both WT mesendoderm (MEN) cells and Foxa2-KO MEN cells^52^. Eo-KO is short for Eomes knockout cells, F2-KO is short for Foxa2 knockout cells. **d.** The experimental flow to determine the biological functions of *Noto* and *Pou6f1*. **e.** UMAP plotting of the acquired single cell multi-omic data for the wildtype (WT), *Pou6f1* KO and *Noto* KO E7.5 embryos. Top: The 10xscMultiome-snRNA-seq data, bottom: 10xscMultiome-snATAC-seq data. **f.** Barplot showing the relative abundance of celltypes in the mutants. **g.** Violin plot showing the AUC score for the down-regulated node-related genes in WT, *Noto* KO and *Pou6f1* KO embryos. **h.** RNAscope analyses validating the expression change of ciliogenesis-related gene-*Foxj1* in *Noto* KO and *Pou6f1* KO embryos. **i. j.** Scan electron microscopy reporting the mean cilia length in the E7.5 mouse embryos of different genotypes (**i**). The wholemount morphologies of embryos were shown at the left with node regions highlighted by white triangles. The zoomed-in views of the node and cilia were shown at the right with white triangles pointing to each cilium. Violin plot showing the statistical summary of cilium length across different genotypes (**j**). Statistical significance were calculated by using Brown-Forsythe and Welch ANOVA tests. **k.** Heatmaps profiling the chromatin accessibilities around the prospective NOTO target peakset (left) and POU6F1 target peakset (right).

To demonstrate this molecular framework, we integrated datasets of both EOMES and FOXA2 ChIP-seq and the chromatin accessibility data of Eomes-KO and Foxa2-KO cells. By profiling the enrichment around the pre-opening *Foxj1-DRE* element (Fig. 5i), we found that the chromatin accessibility for *Foxj1-DRE* element was markedly downregulated in Eomes-KO and Foxa2-KO cells (Extended Data Fig. 12f), which strongly suggests the roles of EOMES and FOXA2 in opening *Foxj1-DRE* chromatin region. We also systematically analyzed the enrichment of EOMES and FOXA2 around the E7.0_APS_M signaling responsive peaks, the NOTO target peaks, and also the POU6F1 target peaks, we found that both EOMES and FOXA2 were enriched around these pre-defined genomic regions (Fig. 6c; Extended Data Fig. 12g, 12h). Moreover, the knockout of either *Eomes* or *Foxa2* led to the reduction of chromatin accessibility around these loci (Fig. 6c; Extended Data Fig. 12g, 12h). Therefore, EOMES and FOXA2 are likely to be responsible for pre-opening signaling responsive elements and NOTO and POU6F1 target chromatin regions.

Next, we investigated whether the TFs-NOTO and POU6F1 play roles in establishing the terminal transcriptomic state but not the chromatin setup. Cross-referencing published atlas^2^ confirmed the enrichment of these two genes in the notochord cells (Extended Data Fig. 13a). We then generated two mouse mutants with the genetic deletions of the *Noto* or *Pou6f1* genes (Fig. 6d; Extended Data Fig. 13b). No visible morphological phenotypes were detected in either mutants at E7.5 stage (Extended Data Fig. 13c). To probe the potential molecular abnormalities, we collected E7.5 WT control, *Noto* KO, and *Pou6f1* KO embryos and performed 10x scMultiome sequencing (snRNA-seq + snATAC-seq). Cells were annotated by computationally mapping their transcriptome onto our E7.5 WT atlas (Fig. 6e). The celltype compositions in the KO embryos remained comparable to WT, except for the axial mesendoderm-related cells, where *Noto* and *Pou6f1* are expressed and now successfully removed (Fig. 6f; Extended Data Fig. 13d, 13e). Moreover, in both *Noto* KO and *Pou6f1* KO embryos, the expression of node cell-related genes and ciliogenesis was severely disrupted (Fig. 6g-j; Extended Data Fig. 13f, 13g; Supplementary Table 5). Phenotypically, directional axis turning, which is related to proper notochord function,^33^ also seem to be randomized (Extended Data Fig. 13h). Importantly, the chromatin accessibility of the TF target chromatin peaks remained unchanged (Fig. 6k; Extended Data Fig. 13i). These results demonstrate that NOTO and POU6F1 function primarily as the final-step transcriptional regulators in the TF relay during the sequential development of the mouse axial mesendoderm.

Together, through systematic exploration of the ST-MAGIC (+) resource, we unraveled the cellular events and associated highly-organized regulatory cascades of TF and signal GRNs, and demonstrate the differential roles for TFs from different stages in shaping gene expression architecture and chromatin accessibility landscape during lineage development (Fig. 7).

**Fig. 7.**
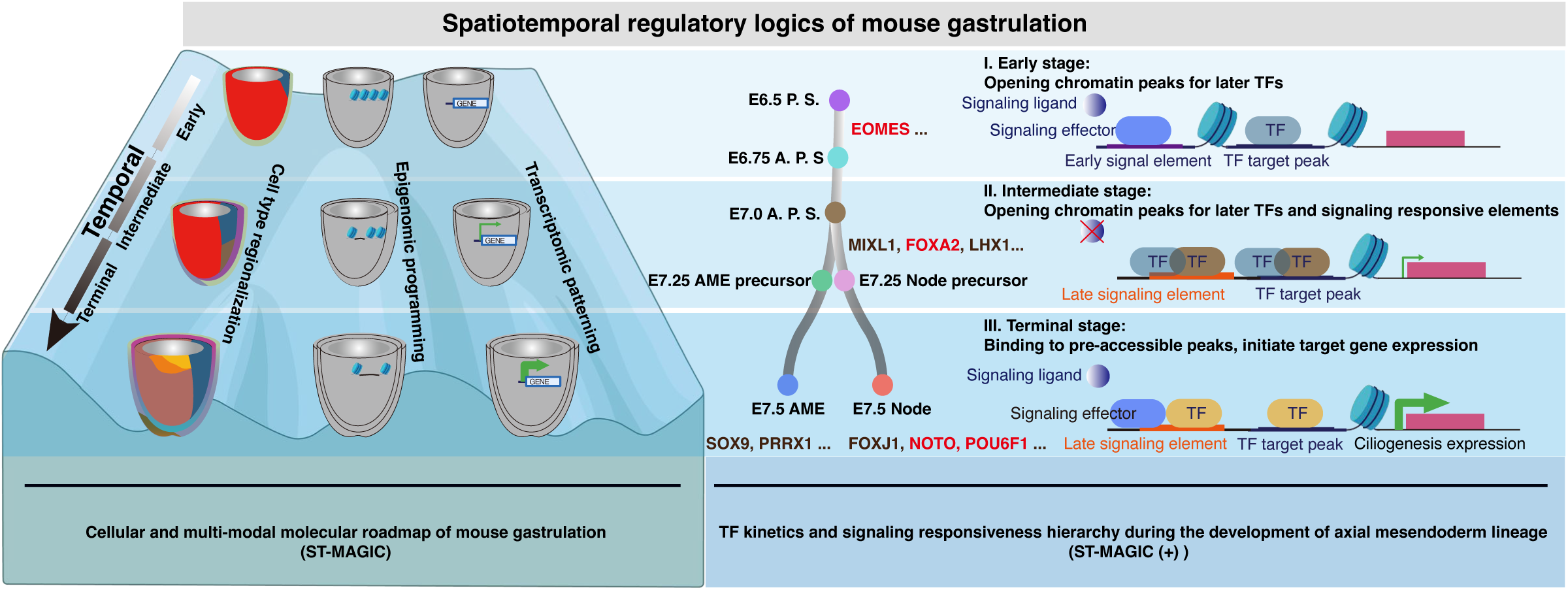
The model summarizing the spatiotemporal logics of cell type regionalization, epigenomic programming and transcriptional patterning during mouse gastrulation and the hierarchy of gene regulatory networks relay underlying axial mesendoderm development.

## Discussion

Gastrulation is a critical developmental stage responsible for generating the three primary germ layers and related derivatives. This process involves scheduling the spatiotemporal sequence of events that leads to the formation and positioning of tissue and organ progenitors.^34^ Understanding the sequential allocation of cell lineages and acquisition of cell fates during gastrulation is challenging, due to the intricate spatiotemporal dynamics involved. In this study, by performing time-series single-cell multi-omic profiling and integrating this with true 3D spatial transcriptome reference data, we established the ST- MAGIC and ST-MAGIC (+) resources, and uncovered an unprecedented level of details of the spatiotemporal GRN dynamics that underlie the sequential cell fate commitment process. This comprehensive integration empowers the exploration of intricate interplays among transcriptomic, epigenomic, and signaling in a spatiotemporal context, providing a detailed view of the molecular landscape during gastrulation.

The refined gene-peak linkage identification method, BioCRE, was developed to address the challenge of linking genes with their regulatory elements. While existing methods can predict regulatory elements, they often lack the gene-peak dual regulation precision required for dynamic processes. BioCRE integrates transcriptomic and chromatin accessibility data to construct robust gene-peak linkages, enabling a more accurate and context-specific understanding of GRNs. Although standard datasets are not yet available to benchmark the efficacy of BioCRE, preliminary results indicate its ability to identify key regulatory elements and their interactions at high precision. This method offers an advantage in uncovering the molecular mechanisms driving cell fate decisions and can be further validated through experimental approaches.

Single-cell omic technologies have significantly broadened the molecular understanding of vertebrate embryogenesis.^4,23,35–38^ Joint analyses of various modal data types, including spatiotemporal dimensions, transcriptome, epigenome, proteome, metabolome, etc, hold promise for achieving a comprehensive understanding of the principles of developmental biology.^39,40^ Cross-modal data integration using bioinformatic strategy is essential to maximize the value of existing data resources. However, integrating different data modalities remains difficult due to the lack of reliable anchors and toolkits. Here, taking transcriptomic information as an anchor to integrate 10x scMultiome with the stage-matched spatial transcriptome coordinates, we constructed the ST-MAGIC atlas with genuine spatiotemporal information, transcriptome architecture, and epigenomic landscape for diverse celltypes in the gastrula. Moreover, by incorporating the eRegulon datasets and published ChIP-seq data of signaling effector, we expanded the ST-MAGIC into ST-MAGIC (+), which enables the delineation of spatiotemporal dynamics of developmental TFs and the characterization of spatiotemporal transition of chromatin responsiveness to developmental signaling during sequential cell fate commitment for all the fated celltypes with unprecedented single-cell resolution in the mouse gastrula.

The ST-MAGIC and ST-MAGIC (+) platforms provide a versatile resource for dissecting the multifaceted mechanisms of embryonic development. Despite the extensive data generated, the scope of this study is limited by the need to focus on specific aspects due to length constraints. We firstly unveiled the multi-omic basis for left-right symmetry breaking event by identifying the potential involvement of distal regulatory elements in regulating the expression of symmetry-breaking genes. We also focused on the axial mesendoderm lineage, particularly the development of the node, to demonstrate the utility of our resource. As known, the axial mesendoderm lineage gives rise to the prechordal plate, anterior head process, and the node-derived notochordal precursors, ultimately forming the notochordal plate on the ventral surface of the mouse embryo.^30,41–43^ We identified the spatiotemporal trajectory of the axial mesendoderm lineage, which originates from cells in the anterior primitive streak and gradually features at the distal tip in E7.5 embryos. By tracing the molecular routes for the two cell subtypes (anterior mesendoderm cells and node cells), we found that both are derived from E7.0 anterior primitive streak cells, when and where the intrinsic chromatin setup for developmental signaling starts to relocate. Furthermore, we identified sequential relays of TF GRNs from early stage, intermediate stage, to terminal stage for each sub-lineage. Through detailed exploration of TF expression, TF-target gene expression, and TF-target peak chromatin accessibility, we characterized the distinct kinetics for early, intermediate, and terminal stage TFs.

Transcription factors emerging at different stages of development may be responsible for distinct functions. Previous studies have revealed genetic deletion of TFs such as *Eomes*^44^ and *Foxa2*^45^ in mouse embryos leads to the complete absence of node structures. Mutation of *Noto* in mouse embryo only shows moderate defects related to ciliogenesis in the node cell, without affecting the formation of the node.^46^ In contrast, the loss of *Noto* homolog, *flh*, in zebrafish embryos leads to the absence of notochord-related cells.^47,48^ These results strongly suggest that the mouse NOTO TF is involved in a lower hierarchy during the notochordal development than EOMES, FOXA2, and even its homolog in zebrafish embryos. Here, as revealed in our study, we found that deletion of early stage TF (EOMES) and intermediate stage TF (FOXA2) severely affects the downstream GRNs, whereas ablation of late stage TFs in mouse embryos only affects the expression of target genes but not the pre-established chromatin setup. Phenotypical characterization showed that knockout mice for terminal stage TFs only leads to the down-regulation of ciliogenesis genes and shortened cilium length. These discrepancies suggest that the post-established chromatin landscape for early and intermediate stage TFs and the pre-established chromatin landscape for terminal stage TFs may build up the molecular basis of developmental competency for node cell formation. Thus, we propose that these distinctions may reflect a molecular ‘priming-specification-determination’ cascade that underlies cell fate commitment. Further exploration of GRN hierarchies across various lineages could define fundamental rules as well as lineage-specific logics that are fit for guiding lineage development during embryogenesis.

Despite the comprehensive nature of our data, several limitations should be acknowledged. First, the current ST-MAGIC and ST-MAGIC (+) resource are limited to spatiotemporal information, transcriptome, and chromatin accessibility at Geo-seq spatial resolution. Transcriptome coordinate profiling using STEREO-seq^12^ or MERFISH^49^, and incorporating additional data types, such as histone modifications, proteomics, and metabolomics, would further enhance the molecular understanding of early embryogenesis. Second, most GRNs for TFs and signaling pathways are defined by integrating consensus targets from public datasets or state-matched in vitro counterparts. Comprehensive profiling of TF or signal targets from precisely matched cell types would improve the reliability of GRN delineation. Third, while the refined BioCRE method provides useful insights into regulatory element and gene-peak linkages, further experimental validation is needed to confirm the predicted regulatory elements and interactions. This includes validating the roles of linked regulatory elements through functional genomics study in embryo models and cell differentiation models.

In summary, this study establishes a comprehensive spatiotemporal multi-omic resource, that recapitulates the sequential cellular events, and dissecting the continuous molecular cascades of TF GRNs and signaling responsiveness during mouse gastrulation. This work opens avenues toward a systematic understanding of the molecular principles governing mammalian embryogenesis in a spatiotemporal context.

## ACKNOWLEDGMENTS

We are grateful to Prof. Chi-Chung Hui for his constructive suggestions. We thank the imaging and animal care core facilities at Guangzhou National Laboratory. We also would like to thank Jiaheng Chen and Heying Li from the core facility at Guangzhou Institutes of Biomedicine and Health, CAS for the help of scan electron microscopy. This work was supported in part by the National Key Basic Research and Development Program of China (2018YFA0800100, 2019YFA0801402), the National Natural Science Foundation of China (32130030, 32470866, 31900454, 32370972), the Major Project of Guangzhou National Laboratory (GZNL2023A02005, GZNL2023A02007, GZNL2023A03005), the Union Project from Guangzhou National Laboratory and State Key Laboratory of Respiratory Disease, Guangzhou Medical University (GZNL2024B01007, GZNL2024B01004), the Guangdong Basic and Applied Basic Research Foundation (2024B1515020052, 2023A1515011783).

## AUTHOR CONTRIBUTIONS

X. Y., S. S. and N. J. conceived the study. X. Y., N. J. designed the experiments, X. Y., P. S. Y. C., Y. Y., Z. L., P. M. collected the sequencing data and performed the experiments, M. W. performed animal husbandry. B. X., S. S. designed the bioinformatic model and pipeline for data analyses and visualization, B. X., C. L., F. T., Y. Y., R. S. performed the bioinformatic analyses. X. Y., B. X., P. T., S. S. and N. J. wrote the manuscript with the help of all other authors.

## DECLARATION OF INTERESTS

The authors declare no conflict of interests.

---

**Extended Data Fig. 1 Quality control analyses of the mouse gastrula single-cell mutli-omic data.**

**a.** Bar-plots showing the quality control metrics of single-cell multiomic (snRNA-seq + snATAC-seq) data. Top panel: number of detected cells, number of detected genes, number of detected UMIs, percentage of mitochondrial RNA for samples detected in transcriptomic data; bottom panel: number of detected cells, number of detected fragments, TSS enrichment scores, blacklist fractions for samples detected in epigenomic data.
**b.** Histograms summarizing the insert size distribution of snATAC-seq fragments (left) and the enrichment of fragments around the nearest gene’s TSS (right) per embryo sample.
**c.** Venn diagram displaying the co-detected cells with two modalities which passed the quality control.
**d.** Bar-plots showing the number of cells captured in this study compared to the number of cells estimated in the corresponding embryonic stage.
**e.** UMAP plot based on 10xscMultiome-snRNA-seq showing the distribution of quality assured cells for each embryonic stage.
**f.** UMAP plot based on 10xscMultiome-snATAC-seq showing the distribution of quality assured cells for each embryonic stage.
**g.** Dynamics of celltype frequency per embryonic stage, the absolute height of stacked celltypes for each stage was scaled according to the predicted cell numbers for corresponding embryonic stage. The dynamic flow shows both the progressive cell number increment and the diversified celltype composition across gastrulation.

**Extended Data Fig. 2 Comparisons between the scMultiome-derived transcriptome and a published dataset.**

**a.** UMAP co-embedding of the scMultiome-derived transcriptome and a published dataset^2^ as marked by data resources (top: scMultiome-derived transcriptome in blue, published dataset in red) and stage information (bottom).
**b.** The bar-plots showing the annotated celltype composition in this study (top) and the published dataset^2^ (bottom).
**c.** The pearson correlation map between celltypes annotated for embryos at each stage in this study and in the published dataset^2^. The right symbol legend was used to represent corresponding celltype annotations. Symbols start with ‘p’ indicates annotations from Pijuan-Sala’s study, symbols start with ‘g’ indicates annotations in this study.
**d.** Dot plot representing the correlation of marker genes (*Sox2*, *Mesp1*, *Sox17*, and *Hnf4a*) expression for celltypes annotated in this study and in the published dataset^2^. Color legend was from Fig. 1b.

**Extended Data Fig. 3 The scMultiome atlas delineates the molecular features of the annotated gastrulating cells.**

**a.** Heatmap showing the differential expressed genes across the 31 annotated celltypes. Representative genes were highlighted at the right.
**b.** Heatmap showing the differential accessible peaks across the 31 annotated celltypes.
**c.** Heatmap showing the TF-motifs enrichment across the 31 annotated celltypes. The representative motifs were highlighted at the right.

**d-f.** Bar plot summarizing the number of linked genes per peak (**d**), the number of linked peaks per gene (**e**), as well as the genomic distance between genes and linked peaks (**f**).

**g.** The enhancer activity reporter assay recording the *in vivo* distribution of enhancer activity of the Neo_ME element across mouse gastrulation. The expression of *Mesp1* detected by wholemount *in situ* hybridization was also shown at the right.
**h.** Top: The diagram showing the experimental workflow for *Mesp1* expression detection during *in vitro* spontaneous embryoid bodies (EB) differentiation. Bottom: Real-time quantitative PCR determining the expression dynamics of *Mesp1* during *in vitro* spontaneous differentiation for wildtype, Neo_ME knockout, and EME knockout cells, respectively. *Mesp1* expression was significantly down-regulated in both Neo_ME knockout and EME knockout cells compared to the wildtype counterpart. The significances (p-value) calculated by two-sides students’ t-test were labeled for day 6 differentiated samples.

**Extended Data Fig. 4 ST-MAGIC captures the spatial evolution of gene expression and chromatin accessibility landscape.**

**a.** Weighted Nearest Neighbor UMAP plotting of multiome dataset from embryonic cells used in ST-MAGIC.
**b.** The Jensen-Shannon divergence (JSD) index evaluating the consistency of gene expression pattern between ST-MAGIC and GEO-seq resource.
**c.** The distribution of single cells mapped to the respective embryonic regions using ST- MAGIC. The density of color dots represents the number of cells mapped to each spot.
**d.** The GEO-seq result (top) and ST-MAGIC visualization (bottom) of *Sox2* expression pattern in the gastrulating embryos..
**e.** RNA scope experiment validating the localization of *Sox2* transcripts in the gastrulating embryos.
**f.** The H3K27ac modified chromatin regions in the E7.5 mouse embryo revealed by ST- MAGIC. Left: the schematic diagram showing the origin of cell samples in the published H3K27ac ChIP-seq data^3^; middle: the representative genome browser snapshot of H3K27ac distribution for selected genomic regions; right: the ST-MAGIC reconstructed corn-plot profile of region-specific H3K27ac modified chromatin regions in the E7.5 mouse embryo.

**Extended Data Fig. 5 ST-MAGIC records the regionalization of gastrulating cell types.**

**a.** ST-MAGIC visualization of celltype composition across the five embryonic stages.
**b. c.** The spatial visualization of major celltypes from the E7.5 mouse embryo in UMAP plot (**b**) and ST-MAGIC (**c**).
**d.** Violin plot showing the expression pattern of marker genes for mesenchyme, cardiac mesoderm, paraxial mesoderm 1, and paraxial mesoderm 2.
**e.** ST-MAGIC visualization of *Hand1*, *Isl1*, *Lefty2*, and *Irx5* expression at the E7.5 mouse embryo.
**f.** ST-MAGIC visualization of chromatin peaks linked to *Otx2* expression at the E6.5 mouse embryo.

**Extended Data Fig. 6 The lateral biased peaks were opened progressively during the gastrulation at E6.75-E7.0.**

**a.** ST-MAGIC visualization of left-right asymmetric gene expression.
**b.** Progressive opening of LPM peaks along with the lineage trajectory of paraxial mesoderm 2 (shown above the heat map) during gastrulation. PS is short for primitive streak cells, the other abbreviations are the same as in Fig. 2k.
**c.** Genome browser snapshot of chromatin accessibilities for LPM related gene, *Lefty2*, and linked peaks.
**d. e.** ST-MAGIC displaying the spatial chromatin accessibility of the *Lefty2* left-right asymmetric specific elements (**d**: the newly identified Neo_LRE element; **e**: the ASE enhancer) during mouse gastrulation.
**f.** Whole mount in situ hybridization showing the expression pattern of *Lefty2* transcripts during gastrulation. The expression of *Lefty2* in the germ layers of E7.25 and E7.5 embryos (at the plane marked by dotted line) were also shown (right panels).
**g.** Violin plot showing the expression of *Lefty2* expression by real-time qPCR analysis in WT and Neo_LRE KO cells during *in vitro* stem cell differentiation. Significances (p-value) were determined by two-sides students’ t-test on day 4 samples.
**h.** Progressive opening of RPM peaks in the lineage trajectory of mesenchyme (shown above the heatmap) during gastrulation.
**i.** Genome browser snapshot depicting the chromatin accessibility landscape and the peak-gene linkage around the *Hand1* locus.
**j.** Visualization of the left-right differences in peak chromatin accessibility of the *Hand1* linked distal peak.

**Extended Data Fig. 7 Identification of specific eRegulon using SCENIC (+).**

**a.** Celltype-specific distribution of the eRegulon specificity score.

**Extended Data Fig. 8 Spatiotemporal reconstruction of eRegulon using ST-MAGIC (+)**

**a.** ST-MAGIC (+) visualization of pre-defined peak sets derived from published dataset^3^.
**b.** Violin plot showing the expression of *Noto* in different celltypes during gastrulation.

**Extended Data Fig. 9 The posterior-enriched intrinsic chromatin responsiveness to WNT signaling involved in mesoderm diversification.**

**a.** The diagram describing the molecular cascade of WNT signaling.
**b.** ST-MAGIC visualization the expression pattern of (i) WNT signaling receptor-*Lrp6* (top) and (ii) effector-*βCatenin* (bottom) during gastrulation.
**c.** Q-Q plot showing the ESC- and EpiLC-specific βCatenin binding peaks.
**d.** Heatmaps showing the βCatenin binding signal in ESC and epiblast-like cells (EpiLCs), with information drawn from dataset of βCatenin ChIP-seq in ESCs^25^ and EpiLCs^26^. Function annotation of the peaks was inferred by GREAT analysis (right panel).
**e.** ST-MAGIC (+) visualization of the chromatin accessible landscape of ESC-specific βCatenin binding peaks (top) and the linked gene expression pattern (bottom) in the gastrulating embryo.
**f.** The 2380 EpiLC-specific βCatenin binding peaks and the 801 linked genes were reconstructed to determine the spatiotemporal dynamics of chromatin responsiveness (top) and transcriptional output of WNT signaling (bottom).
**g.** The model summarizing the morphogen distribution (top), intrinsic chromatin responsiveness (middle) and transcription output (bottom) of WNT signaling inferred from ST-MAGIC (+) results.

**Extended Data Fig. 10 The transition of chromatin responsiveness to NODAL signaling during gastrulation.**

**a.** The diagram describing the molecular cascade of NODAL signaling.
**b.** ST-MAGIC visualization the expression pattern of NODAL signaling receptors-*Alk4* and *ActRIIB* (top) and effectors-*Smad2* and *Smad3* (bottom) in the gastrula.
**c.** ST-MAGIC visualization of the expression pattern of NODAL signaling antagonist-*Cer1* during gastrulation.
**d.** Q-Q plot showing the ESC- and EB-specific Smad2/3 binding peaks.
**e.** Heatmaps showing the NODAL signal effector-Smad2/3 binding signal at both pluripotent state cells (ESCs) and lineage primed state cells (EBs). The published dataset for Smad2/3 ChIP-seq from ESCs and EBs^29^ were used. The biological function annotation of the peaks using GREAT analysis was shown on the right.
**f.** The model summarizing the morphogen distribution (top), intrinsic chromatin responsiveness (middle) and transcription output (bottom) of NODAL signaling inferred from ST-MAGIC (+) results. The spatiotemporal shift of NODAL signaling responsiveness was clearly captured.
**g.** GREAT analysis (biological processes enrichment (left), mouse phenotype enrichment (right)) of the E7.0_APS_M NODAL signaling responsive peaks.

**Extended Data Fig. 11 The emergence of node and anterior mesendoderm in E7.5 mouse gastrula.**

**a.** The celltype composition at the E7.5 distal tip region highlighting the enrichment of node and anterior mesendoderm cells. Right: The relative frequencies of corresponding celltypes.
**b.** Top: UMAP plot showing the expression of *Shh*, *Noto*, *Foxj1* in the annotated node and anterior mesendoderm cells. Bottom: Violin plot showing the expression of *Shh*, *Noto*, *Foxj1* in node and anterior mesendoderm cells.
**c.** Wholemount RNAscope staining of *Shh*, *Noto*, *Foxj1* revealing the existence of two celltypes during axial mesendoderm to notochordal development.
**d.** Gene ontology annotation of differentially expressed genes between node and anterior mesendoderm.

**Extended Data Fig. 12 The identification of the intricate molecular relay during axial mesendoderm development.**

**a.** Line plot describing the TF expression, TF-target gene expression, as well as TF-target peak accessibility of Mixl1 (top) and Lhx1 (bottom) during axial mesendoderm lineage.

**b-e.** ST-MAGIC (+) visualization of the gene regulatory network for early stage TF-Eomes (**b**), intermediate stage TFs-Mixl11 (**c**), Foxa2 (**d**), and Lhx1 (**e**) during gastrulation.

**f.** The representative genome browser snapshot reporting the binding enrichment of Smad2/3, EOMES, and FOXA2 around *Foxj1* locus.
**g. h.** Heatmaps showing the enrichment of early stage and intermediate stage TFs on E7.0_APS_M WNT signaling responsive peaks (**g**) and E7.0_APS_M NODAL signaling responsive peaks (**h**) and the dynamics of chromatin accessibilities following the deletion of EOMES or FOXA2. Published dataset of ATAC-seq for both WT EB cells and Eomes-KO EB cells^51^, ATAC-seq for both WT MEN cells and Foxa2-KO MEN cells^52^ were used.

**Extended Data Fig. 13 Expression of node cells marker genes were severely downregulated while the epigenome remained unchanged in terminal stage factors KO mouse models.**

**a.** Violin plot showing the expression of *Noto* (left) and *Pou6f1* (right) from https://marionilab.cruk.cam.ac.uk/MouseGastrulation2018/.
**b.** The genome snapshot for *Noto* (left) and *Pou6f1* (right) genes. The scissors represent the exact cutting sites for genome deletion of these two genes.
**c.** The global morphologies remain no significant aberrations for the *Noto* KO and *Pou6f1* KO embryos at the E7.5 stage.
**d.** Gene expression profiling for both *Noto* (top) and *Pou6f1* (bottom) among WT, *Noto* KO and *Pou6f1* KO samples. Data from the 10xscMultiome-snRNA-seq dataset for the embryos. The expression of *Noto* and *Pou6f1* is absent in the respective KO mutant embryos consistent with the deletion of the genes.
**e.** Genome browser snapshot around the *Noto* (left) and *Pou6f1* (right) loci reporting the complete deletion of the relative genomic sequence in *Noto* KO and *Pou6f1* KO embryos, respectively.
**f.** Bar plot showing the gene ontology analyses for down-regulated node-related genes (related to Fig. 6g).
**g.** Violin plot revealing the expression change of *Foxj1* in *Noto* KO and *Pou6f1* KO embryos.
**h.** The randomization of direction of axis turning in *Noto* KO and *Pou6f1* KO embryos.
**i.** Genome browser snapshot showing no change in chromatin accessibility of the *Foxj1* linked distal peaks in *Noto* KO and *Pou6f1* KO embryos.

**Supplementary Table 1. Celltype-specific genes and linked peaks.**

**Supplementary Table 2. Top1000 specific peaks for RPM and LPM.**

**Supplementary Table 3. Spatiotemporal turnover of WNT signaling responsiveness element.**

**Supplementary Table 4. Spatiotemporal turnover of NODAL signaling responsiveness element.**

**Supplementary Table 5. Downregulated Node cell markers in Noto KO and Pou6f1 KO embryo.**

**Supplementary Table 6. Oligos used in this study.**

## METHODS

### Ethics statement

Mouse used in this study were housed and bred in the SPF facilities of Guangzhou National Laboratory. All animal experiments were performed in compliance with the guidelines of the Animal Core Facility.

### Mouse embryo collection and multi-omic profiling

For embryo sampling, C57BL/6J embryos were harvested from pregnant mice at day 6.5, 6.75, 7.0, 7.25, and 7.5 of gestation (day of vaginal plug detection = Day E0.5). Plugged female mice were picked after mating and marked as embryonic day 0.5 (E0.5). Female mice were sacrificed for embryos collection at specific gestational stages. Embryos were isolated from the uterus and carefully transferred into pre-cool PBS in petri dishes, and surrounding decidua and parietal endoderm tissues were removed using needles under Olympus stereoscope.

Careful morphological staging of the acquired embryos was performed before single-cell multi-omic data preparation. The developmental time points of embryos were staged by the proximal-distal span of the PS and the anterior-posterior span of the mesoderm layer. Embryos from the same stage were pooled together and subjected to TrypLE Express enzyme (Gibco, 12604) incubation at 37°C for 7-10 min. The acquired single-cell suspension was carefully washed and filtered to ensure proper integrity and avoid cell clumping of each single cell. Cell counts were then assessed with a haemocytometer counted under a microscope. Nuclei isolation and multiome library preparation were performed by following the manufacturer’s instructions (https://www.10xgenomics.com/cn/support/single-cell-multiome-atac-plus-gene-expression/documentation).

For the single-cell multi-omic profiling of the Noto KO and Pou6f1 KO embryos, parents of heterozygotic mutation were crossed and checked for the vaginal plug. E7.5 embryos were freshly harvested and a tiny portion of extraembryonic tissues were collected and genotyped. During the genotyping process, the embryos were freshly frozen in liquid nitrogen. Embryos with the same genotypes (WT, Noto KO homozygote, Pou6f1 KO homozygote) were grouped and dissociated into single nuclei. ∼15,000 nuclei for each group were collected and loaded for further single-cell multiome library preparation following the manufacturer’s instructions.

### Enhancer activity reporter assay

Generally, the enhancer reporter assay was performed as previously reported.^3^ In brief, DNA fragments for potential regulatory elements were cloned from the C57BL/6J mouse genome and then ligated into the plasmid construct containing the minimal Hsp68 promoter and LacZ. The acquired purified plasmids were then linearized and used for pronuclear injections of PN4 stage zygotes with a FemtoJet Microinjection System (Eppendorf). The injected embryos were cultured to the 2-cell stage in KSOM medium with amino acids at 37 °C under 5% CO2, and then transferred to the oviduct of pseudo-pregnant ICR females and marked as 0.5 dpc. Embryos were collected at the corresponding stage for LacZ staining. LacZ staining was performed using a commercialized β-gal staining kit (Beyotime, RG0039).

### RNAscope staining for embryo whole mount or sections

RNAscope probes including mm-Lefty2-C2 (436291-C2), mm-Hand1-C1 (429651), mm-Irx5-C2 (513871-C2), mm-Isl1-C3 (451931-C3), mm-Sox2-C1 (401041-C1), mm-Otx2-C3 (444381-C3), mm-T-C3 (423511-C3), mm-Eomes-C2 (429641-C2), mm-Foxa2-C4 (409111-C4), mm-Shh-C2 (314361-C2), mm-Noto-C3 (1253281-C3), mm-Foxj1-C3 (317091-C3), mm-Dynlrb2-C3 (1243011-C3), mm-Col2a1-C4 (407221-C4), mm-Rsph9-C2 (430201-C2), and mm-Pou6f1-C1 (801931-C1) were bought from the Advanced Cell Diagnostics.

For RNAscope staining with wholemount embryos, embryos were carefully fixed in 4% PFA overnight. After serial dehydration and rehydration of embryos using gradient methanol-PBS solution, embryos were subject to RNAscope protocol following the manufacturer’s instruction(https://acdbio.com/sites/default/files/MK%2050016%20Zebrafish_WISH_Tech%20Note_12042017.pdf). The final images were acquired through using the LiTone XL system (Light Innovation Technology Limited).

For RNAscope staining with embryo sections, dissected embryos were first fixed in 4% PFA, then dehydrated in 20% Sucrose-PBS and 30% Surcrose-PBS solution, respectively. The dehydrated embryos were then subjected to OCT embedding before cryo-sectioning using Leica CM1950. The following RNAscope workflow was performed by following the manufacturer’s instructions (https://acdbio.com/ebook/introduction/materialsmethod). Images were finally collected by using the Carl Zeiss LSM980 system.

### Wholemount in situ hybridization

Wholemount in situ hybridization of RNA transcripts was performed by following the published protocol.^53^ Briefly, DNA fragments encoding the probes of *Lety2* were firstly PCR amplified by using the oligos in Supplementary Table S6 using a mouse embryo cDNA library. Embryos at relevant stages were collected in DMEM media and then fixed in 4% PFA at 4°C overnight. Fixed embryos were then washed in PBS solution at room temperature to remove residual PFA, followed by serial dehydration and rehydration in PBS, 25% Methanol/PBS, 50% Methanol/PBS, 75% Methanol/PBS, and 100% Methanol. Afterward, the embryos were treated with 10 μg/mL proteinase K and incubated with DIG-labeled RNA probes at 70°C overnight. To remove the remaining RNA probes, embryos were washed in TBST buffer with frequent buffer changing for at least 2-4 hours. Then the embryos were incubated with anti-DIG-AP antibody at 4°C overnight. The embryos were subjected to sufficient TBST wash before final staining with NBT and BCIP solution. The final images of stained embryos were collected using an Olympus SZX16 microscope.

### The generation of genome edited mouse embryonic stem cell or mouse model

In this study, we have prepared the genome-edited mouse embryonic stem cell lines with *Mesp1* Neo_ME element, *Mesp1* EME element, and *Lefty2* Neo_LRE element knockout, respectively. CRISPR-Cas9 system was used to generate the genome-edited mouse embryonic stem cell lines. Small guided RNAs were designed by using the online tool Chop-chop (http://chopchop.cbu.uib.no/). The synthesized sgRNA DNA fragments were then ligated into the px330 plasmid with a Cas9 protein expression cassette. The acquired purified plasmids were then transfected into the WT cells using Lipofectamine 3000. The genome-edited clones were then selected and genotyped as previously reported^54^. The potential off-target editing sites were tested based on the website’s indication.

To determine the roles of *Pou6f1*, we also generated a mouse model with *Pou6f1* deletion. The specific sequences for sgRNA pairs were included in Supplementary Table S6. DNA fragments for the sgRNA and T7-Cas9 were transcribed and purified *in vitro* using MMESSAGE MMACHINE T7 Ultra Kit (Invitrogen, AM1345) and MEGAclear kit (Invitrogen, AM1908). To prepare a sufficient number of fertilized zygotes, C57BL/6J female mice (4 weeks old) were superovulated and mated with the male C57BL/6 mice. Twenty-four hours later, fertilized embryos were collected from oviducts. RNA for Cas9 protein (100 ng/µl) and corresponding sgRNA (100 ng/µl) were mixed in HEPES-CZB medium containing 5 μg/ml cytochalasin B (CB) and injected into the cytoplasm of fertilized eggs using a FemtoJet microinjector (Eppendorf) with constant flow settings. The injected embryos were cultured in KSOM with amino acids at 37 °C under 5% CO2 in the air to reach the 2-cell stage after 24 h *in vitro.* Two-cell embryos were transferred into pseudo-pregnant ICR female mice. The acquired mouse individuals were then subjected to genotyping for successful Pou6f1 deletion. Mice with Pou6f1 deletion were retained and recorded as F0. To prepare the stable Pou6f1 deletion mouse line, F0 mice were crossed and the acquired F1 mice were checked. Pou6f1 KO heterozygotes F1 mice were maintained and the population was expanded for the following experiments.

The Noto KO mouse line was directly bought from GemPharmatech (Strain ID: T017401).

### Scanning Electron Microscopy

Embryo samples were freshly collected and fixed in 2.5% glutaraldehyde overnight at 4°C. Subsequently, the samples were thoroughly washed with PBS four times at 10-minute intervals to ensure the complete removal of the fixative. Fixed embryos were treated with osmium tetroxide for 1 hour to enhance the contrast and stability of cellular structures under the electron beam. The samples were then washed with ddH2O five times at 10- minute intervals to remove any residual osmium tetroxide. Dehydrate the embryos in an ethanol series for 5-10 min each: 50%, 70%, 85%, and three times in 100%. Transferred the embryos in ethanol to baskets for critical point drying (CPD) in a critical point dryer machine (Quorum K850 critical point dryer). After that, the samples were mounted on sample stubs and sputter-coated with gold using a Quorum Q150R S sputter coater. Finally, the samples were imaged using the Zeiss GeminiSEM 300 scanning electron microscope.

## Notes

### Competing Interest Statement

The authors have declared no competing interest.

